# Epigenetic regulation by TET1 in gene-environmental interactions influencing susceptibility to congenital malformations

**DOI:** 10.1101/2024.02.21.581196

**Authors:** Bernard K. van der Veer, Lehua Chen, Spyridon Champeris Tsaniras, Wannes Brangers, Qiuying Chen, Mariana Schroiff, Colin Custers, Harm H.M. Kwak, Rita Khoueiry, Robert Cabrera, Steven S. Gross, Richard H. Finnell, Yunping Lei, Kian Peng Koh

## Abstract

The etiology of neural tube defects (NTDs) involves complex gene-environmental interactions. Folic acid (FA) prevents NTDs, but the mechanisms remain poorly understood and at least 30% of human NTDs resist the beneficial effects of FA supplementation. Here, we identify the DNA demethylase TET1 as a nexus of folate-dependent one-carbon metabolism and genetic risk factors post-neural tube closure. We determine that cranial NTDs in *Tet1*^-/-^ embryos occur at two to three times higher penetrance in genetically heterogeneous than in homogeneous genetic backgrounds, suggesting a strong impact of genetic modifiers on phenotypic expression. Quantitative trait locus mapping identified a strong NTD risk locus in the 129S6 strain, which harbors missense and modifier variants at genes implicated in intracellular endocytic trafficking and developmental signaling. NTDs across *Tet1*^-/-^ strains are resistant to FA supplementation. However, both excess and depleted maternal FA diets modify the impact of *Tet1* loss on offspring DNA methylation primarily at neurodevelopmental loci. FA deficiency reveals susceptibility to NTD and other structural brain defects due to haploinsufficiency of *Tet1*. In contrast, excess FA in *Tet1*^-/-^ embryos drives promoter DNA hypermethylation and reduced expression of multiple membrane solute transporters, including a FA transporter, accompanied by loss of phospholipid metabolites. Overall, our study unravels interactions between modified maternal FA status, *Tet1* gene dosage and genetic backgrounds that impact neurotransmitter functions, cellular methylation and individual susceptibilities to congenital malformations, further implicating that epigenetic dysregulation may underlie NTDs resistant to FA supplementation.

## Introduction

Neural tube defects (NTDs) are among the most common congenital malformations, affecting 0.3- 200 per 10,000 pregnancies globally with regional variation in incidences(*1*). They are caused by a failure of the neurulation process early in embryogenesis, when the neural plate folds into a tube to form the morphological scaffold for the developing central nervous system (*2–4*). Failure of neural tube closure (NTC) at the cranial and spinal regions results in the most prevalent open NTD subtypes- anencephaly and spina bifida, respectively (*5*). Folic acid (FA) supplementation, when taken at 0.4 to 0.8 mg daily before or early during pregnancy, has a substantial net benefit to prevent NTDs, but 30-50% of cases remain non-responsive to FA (*6*). How individual genetic differences can result in varied responsiveness to FA supplementation is largely unknown. Despite aggressive campaigns for FA supplementation and fortification of the US food supply, NTDs still affect an estimated 3000 pregnancies in the US annually and remain a significant global public health concern (*7, 8*).

The etiology of NTDs is currently postulated to be multifactorial with contributions from both genetic and environmental factors (*9–11*). In mouse models, NTDs are most frequently observed as exencephaly, the developmental precursor of anencephaly (*5, 12*). Extensive investigations in mice have identified over 300 genes which, when mutated, result in NTDs (*13*). Of these, only some have been orthogonally validated in human genome sequence analysis of relatively small patient case numbers, including many in the planar cell polarity (PCP) (*14–17*) and WNT signaling pathways (*18*). Nonetheless, NTD phenotypes in single gene knockout (KO) mouse models are often incompletely penetrant, suggesting the involvement of multiple genes in the disease (*12, 19*). In the human population, the risk of re-occurrence and pattern of inheritance of NTDs do not follow Mendelian rule, consistent with a polygenic etiology that can involve compound effects of heterozygous mutations and further influences by environmental factors (*20*). While the mouse remains the most common animal model to study the role of candidate genes in the development of NTDs, the contribution of genetic heterogeneity in strain backgrounds is rarely investigated (*13*).

As the largest modifier of NTD risk, folate is a key co-factor in a network of interlinked metabolic reactions that provide one-carbon (1C)-units for nucleotide biosynthesis (*21*). The folate cycle feeds into the methionine cycle to generate S-adenosylmethionine (SAM), a universal methyl-donor required for methylation of DNA, RNA and proteins. Human and animal studies suggest that folate status influence global patterns of DNA methylation, which occurs predominantly at CpG dinucleotides in mammalian DNA (*22–24*). Maternal folate deficiency may increase the risk of NTD by inducing DNA hypomethylation (*25–28*). Mice deficient in DNA methyltransferase 3A (DNMT3A) and DNMT3B, which are responsible for *de novo* re-methylation of the majority of the embryonic genome during peri-implantation development, had an increased risk of NTDs and other embryonic malformations, indicating that impaired DNA methylation can disrupt NTC (*29*).

The role of DNA demethylation during early embryogenesis is more recently understood with the discovery of the Ten-Eleven-Translocation (TET1, TET2, and TET3) DNA dioxygenases. The catalytic activity of TET proteins on DNA results in reiterative oxidation of 5mC into 5- hydroxymethylcytosine (5hmC) and further oxidation products which are subsequently replaced with unmethylated cytosines, either passively by DNA replication or actively by base excision repair (*30–32*). We have previously demonstrated a dominant role of TET1 in the mouse embryo early post-implantation, involving both catalytic and non-catalytic activities in lineage gene regulation (*33*). While high *Tet1* expression is sustained in the epiblast until the onset of gastrulation, loss of function of TET1 results in both repressive DNA and histone hypermethylation that persist in post-gastrulation neuroepithelial cells (*34*). Intriguingly, the penetrance of embryonic lethality and cranial NTDs in *Tet1* null mice are strongly influenced by genetic background, being low in inbred congenic strains (∼20%) but nearly complete (∼80%) in outbred stocks (*33, 35*). This led us to hypothesize that TET1 may be an epigenetic regulator at the nexus of gene-environmental interactions, particularly involving folate 1C metabolism, that profoundly influence susceptibility for congenital malformations including NTDs.

In this study, we backcrossed *Tet1* mutant mice from the C57BL/6J (B6) background to an alternative 129S strain to create an incipient congenic strain that exhibit increased susceptibility to develop NTDs. Using NTD as a trait, we performed strain intercrosses between NTD-resistant (B6129S6F1) and susceptible (129S6.B6) strains of *Tet1* mutants and identified a strong quantitative trait locus (QTL) on chromosome 9 centered at *Csnk1g1*, *Snx1,* and *Dapk2* as the top candidate modifier genes. We characterized wild-type (WT) and *Tet1* null mouse embryonic stem cells (ESCs) derived *in vitro* from B6129S6F1 and 129S6.B6 strains and observed a cell non-autonomous effect of 129S6.B6-*Tet1* null cells to induce NTDs *in vivo* in chimeric embryos. During *in vitro* differentiation of ESCs to neuronal progenitors, *Tet1*-deficient cells exhibit strain differences in sensitivity towards Wnt and Nodal signaling. By examining post-NTC mouse embryos exposed to excess or depleted maternal dietary FA in different strains, we determined *Tet1* gene dosage interactions with folate status underlying NTD susceptibility, resistance to FA supplementation, and differential DNA methylation that converge on the regulation of folate uptake, phospholipid metabolism and neurotransmitter functions. These studies reveal *Tet1* as a pivotal epigenetic connector of developmental signaling, folate one-carbon metabolism and genetic risk factors in gene-gene and gene-environmental interactions that influence birth defect susceptibilities.

## Results

### Loss of *Tet1* results in neural tube defects with varying strain-dependent penetrance

In our previous work, we described a pivotal role of TET1 in orchestrating cell lineage choice during the onset of gastrulation in mouse embryos. The absence of TET1 function resulted in post-gastrulation embryonic defects (*33, 36, 37*). In a *Tet1*^GT(RRG140)^ strain expressing the TET1 exon2-β- geo fusion protein, we initially observed embryonic lethality with varying penetrance, dependent on the genetic background. It was fully penetrant in mixed B6;129P2/OlaHsd, became sub-penetrant with rescue of viability after 6 generations of backcrossing to C57Bl/6J (B6) but resumed 100% lethality upon outbreeding to CD1 (*33*). To eliminate the possibility of embryonic lethality due to dominant negative effects of ectopic gene trap (GT) fusion protein expression in *Tet1*^GT/GT^ mice, we engineered a new gene-targeted B6-*Tet1*^tm1Koh^ mouse strain, in which an insertion cassette at the first coding ATG effectively abolishes all coding transcripts of full-length *Tet1* expressing an embryonic isoform (*33*). In *Tet1*^-/-^ (KO) embryos of the B6-*Tet1*^tm1Koh^ strain, we observed a 25% failure in anterior neuropore closure without developmental delay or other gross abnormalities, indicating a specific impact on cranial NTC. As in other *Tet1*^-/-^ models, homozygous loss of *Tet1* in the B6 background results in restricted growth and subfertility in mice, despite post-natal viability (*33, 35, 38*).

To demonstrate strain-dependency in the penetrance of the NTD phenotype in *Tet1* KO, we outcrossed the B6-*Tet1*^tm1Koh^ congenic strain to genetically heterogeneous outbred CD1 lines for at least three generations. Indeed, NTDs specific to *Tet1*^-/-^ embryos were observed at rates >70%, confirming a significant contribution of genetic modifiers to the phenotype, similar to *Tet1*^GT/GT^ mice (Fig. 1A). Further characterizing the NTDs in CD1-*Tet1* KO embryos from E9.5 until E12.5, we observed a progression from failed closure of the E9.5 cranial neural tube to exencephaly, a protrusion of brain tissue outside of the cranial vault by E11.5 (*5*) (Fig. 1B). Histologically, we confirmed incomplete anterior midbrain-hindbrain neuropore closure at closure site 2 in *Tet1*^-/-^ embryos, while the posterior neuropore closed completely by E10.5 (Fig. 1C). No overt differences in Theiler stage or somite counts were observed between NTD-affected and normal E11.5 embryos (data not shown). At E11.5, *Tet1*^+/+^, *Tet1*^+/-^ and *Tet1*^-/-^ embryos were obtained from heterozygote intercrosses in the expected Mendelian ratio in all genetic backgrounds (Supplementary Fig. 1A), suggesting there was no attrition of KO embryos due to earlier embryonic lethality.

**Fig. 1.**
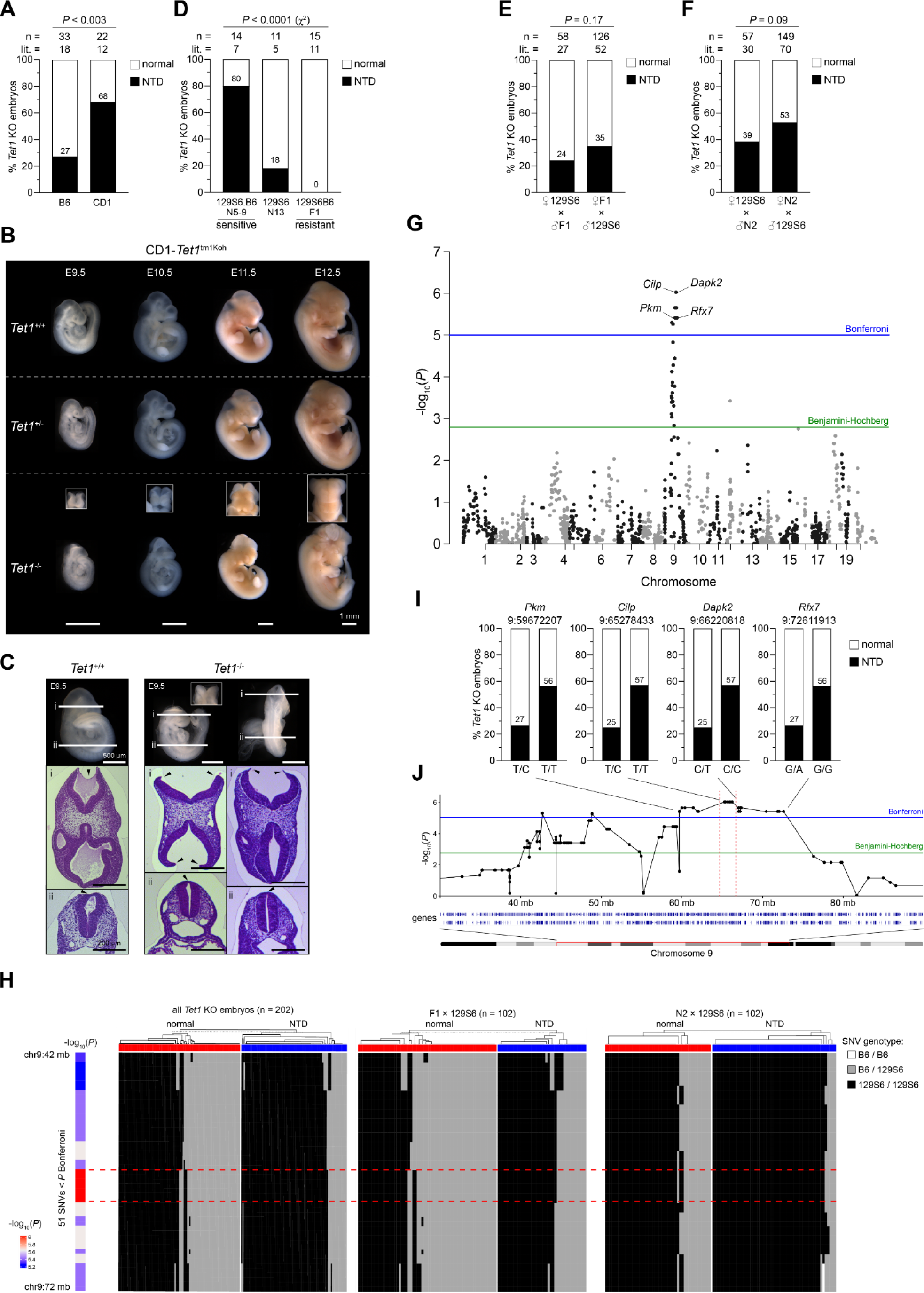
Phenotypic analysis of NTD-resistant and -susceptible strains of Tet1-/- mice and QTL mapping to a chromosome 9 risk locus. (**A**), NTD rates of *Tet1*^-/-^ embryos in B6 and CD1 genetic backgrounds. n = number of embryos, litter (lit.) = number of litters collected. The exact percentage of NTD is indicated in each column. *P* values are calculated by Fisher’s Exact test. (**B**), Images of E9.5 to E11.5 embryos per genotype of CD1-*Tet1*^tm1Koh^ mice. In *Tet1*^-/-^ embryos, the failure of NTC is shown with a frontal view image (above) of the forebrain at each stage. Scale bar indicates 1 mm. (**C**), Hematoxylin & eosin-stained sections of E9.5 CD1-*Tet1*^+/+^ and *Tet1*^-/-^ embryos. Lines in whole embryo image (top) indicate planes of section at cranial (i) and caudal (ii) regions. Arrowheads indicate failed neuropore closure in cranial (i) but normal closure in caudal (ii) sections of *Tet1*^-/-^ embryos. Scale bar indicates 200 µm. (**D**), NTD rates of *Tet1*^-/-^ embryos upon backcrossing of B6 to 129S6 strain. Overall significance in strain-dependent differences is determined using Chi square test. (**E**) and (**F**), NTD rates of *Tet1*^-/-^ offspring of heterozygous parents in reciprocal intercrosses (maternal x paternal) between F1 (E) or N2 (F) generation and 129S6 strains. *P* values are calculated by Fisher Exact test. (**G**), Manhattan plot representing genome-wide association of whole-exome sequencing single nucleotide variants (n = 5544) with NTD phenotype in 103 embryos from N2 x 129S6 and F1 x 129S6 crosses. Each point represents a SNV. The horizontal blue line corresponds to a genome-wide significance threshold (*P*<1x10^-5^) based on Bonferroni correction, and the green line corresponds to a threshold (*P*<1x10^-3^) based on Benjamini-Hochberg correction (FDR). (**H**), Heatmap representation of the genotype of 51 SNVs (Bonferroni corrected p-value < 0.05) in rank order (rows) in every embryo classified by phenotype (column). Left, all *Tet1*^-/-^ embryos sequenced; middle, F1 x 129S6; right, N2 x 129S6. The 7 SNVs with the max *P*-value are highlighted with red dashed lines or bars in H and J. (**I**), Allele effect plots of SNVs within the most significant QTL peak region. (**J**), Line plot of genotype-phenotype association within a 60 megabase (MB) region on chromosome 9 encompassing the QTL peak. SNVs shown in h are indicated by connecting lines.

We investigated whether the high penetrance of NTDs in *Tet1* KO embryos depends on genetic heterogeneity present in outbred or mixed strains, or could be recapitulated in an alternative inbred congenic strain. To explore this, we backcrossed the B6-*Tet1*^tm1Koh^ allele to an available 129 substrain (129S6, also known as 129/SvEvTac). Interestingly, incipient congenic 129S6.B6-*Tet1*^-/-^ embryos (within 5-9 backcross generations) exhibited a peak in phenotype penetrance as high as 80%. Further backcrossing (≥ 13 backcross generations) to create a highly inbred 129S6 congenic (129S6.Cg) strain reduced NTD rates in *Tet1*^-/-^ to below 20%, suggesting a contribution of both B6- and 129S6-specific variants to NTDs in *Tet1^-/-^* embryos. Intercrossing *Tet1*^tm1Koh^ heterozygous (HET) B6 and 129S6 congenic mice to create F1 hybrids yielded KO embryos completely resistant to NTDs (Fig. 1D), indicating that the NTD causal variants act recessively. These observations suggest that the phenotypic expression of NTDs in *Tet1*-deficient embryos may involve polygenic interactions, such as combinations of heterozygous and homozygous digenic interactions, occurring more frequently in a genetically diverse background than in conventional highly inbred strains.

### QTL analysis maps an NTD risk locus on chromosome 9

The stark variation in NTD penetrance caused by *Tet1* KO across diverse genetic backgrounds suggests the presence of genetic modifiers of large effect size. The high NTD penetrance in incipient congenic 129S6.B6-*Tet1*^-/-^ embryos further indicates that the 129S6 genome carries a majority of NTD risk alleles. To map the genomic location of the genetic modifier(s) through quantitative trait locus (QTL) analysis, we initiated HET x HET intercrosses between the NTD-resistant (0% NTD) F1-*Tet1*^tm1^ and the NTD-susceptible (80% NTD) 129S6.B6 incipient congenic strains. Intriguingly, NTD rates dropped by more than 50% (to about 30%) in the KOs in N2 offspring, even though half were expected to retain the 129S6.B6 genome (Fig. 1E). An additional generation of backcrossing N2 HET mice to the 129S6 strain was required to produce *Tet1*^-/-^ offspring with ∼40-50% NTDs, suggesting the involvement of more than one or two risk alleles (Fig. 1F). To address loss of heterozygosity among informative single nucleotide variants (SNVs) in the N2 progeny, we randomized and limited the use of N2 males to no more than two productive timed mating. The observed genotypes of the offspring were Mendelian, and phenotypes specific to *Tet1*^-/-^ embryos were binary – either exencephalic or normal (Supplementary Fig. 1B-C). In total, 390 *Tet1*^-/-^ embryos were collected, of which 231 classified as normal (controls) and 159 as affected (cases with exencephaly), yielding an overall penetrance of ∼40% from 224 timed mating. When strain crosses were classified by maternal x paternal mating, slightly higher NTD rates were observed in the offspring from F1 x 129S6.B6 and N2 x 129S6 compared to the reciprocal 129S6 x F1 and 129S6 x N2 mating, respectively (Fig. 1E-F). Moreover, a slight female excess in KO embryos displaying NTDs was notable (Supplementary Fig. 1D-F). However, none of these sex-linked biases reached statistical significance, suggesting that any maternal contribution of genetic variation to the phenotype is likely to be weak.

Proceeding to the QTL analysis, we conducted exome sequencing of a subset of 208 KO embryos from F1 x 129S6 and N2 x 129S6 crosses, of which 202 passed quality control for genome-wide genotype-phenotype association using the NTD phenotype as a trait. Among 5544 SNVs detected over all chromosomes, 51 SNVs surrounding a chromosome 9 (chr9) locus surpassed significance thresholds after a stringent Bonferroni correction. The number of significant SNVs at the locus expanded to 217 using a less stringent threshold based on Benjamini-Hochberg (FDR) correction. Of these 217 less stringent SNVs, all but one map to a 35 megabase (Mb) region (mm10 chr9:40 Mb-75 Mb) surrounding the chr 9 QTL containing 370 protein-coding genes, confirming the presence of one dominant risk modifier locus (Fig. 1G). Charting the genotype of the 51 stringent SNV hits across all 202 embryo samples revealed that 43 highly significant SNVs segregated together at the chr9:57 Mb-76 Mb region (Fig. 1H). Overall, 56-57% of *Tet1*^-/-^ embryos that were homozygous for the 129S6 variants across the QTL developed NTDs, compared to 25-27% in those that were heterozygous (Fig. 1I,J). Nonetheless, we observed sufficient heterogeneity in genotypes between control and NTD-affected embryos to distinguish a 0.9 Mb peak region within the QTL harboring seven SNVs with *p*-values of highest significance, associated with five unique genes between *Cilp* and *Dapk2* (Fig. 1H,J). SNaPshot multiplex PCR genotyping of five of these SNVs at the *Cilp*-*Dapk2* locus in an additional cohort of 141 samples (F1 x 129S6, 14; N2 x 129S6, 20; 129S6 x F1, 53; 129S6 x N2, 54) did not reduce the resolution of the 1 Mb interval but validated this QTL (Supplementary Fig. 1G,H). The incomplete penetrance (50% overall) of NTDs in mice homozygous for these 129S6 risk variants suggests the presence of other QTLs not identified by our genetic crosses involving only two strains. Nonetheless, the identification of a dominant risk allele affirms that a genetic interaction between *Tet1* and modifier gene(s) influences susceptibility to NTDs.

### The *Cnsk1g1*-*Snx1-Dapk2* gene cluster is located centrally at the QTL peak

Since genes in topology associated domains (TADs) are often co-regulated, we assessed the higher chromatin structure over the QTL to compartmentalize SNVs and their associated genes within contact domains. Using publicly available HiC chromatin interaction datasets generated in mouse embryonic stem cells (ESCs), neuroprogenitor cells (NPCs) and cortical neurons (CN) (*39*), we charted TADs over the chr9 QTL (*40*). Interestingly, the 35 Mb stretch (chr9:40 Mb-75 Mb) encompassing all 216 significant SNVs overlapped with a dominant TAD associated with active chromatin, or A-compartment, in all three cell types, suggesting that these loci are active throughout development (Supplementary Fig. 2A). To further select active genes within the TAD, we used RNA-seq datasets of mouse E8.5-10.5 embryonic tissues from the Expression Atlas (*41*). We filtered for active genes based on detectable normalized transcript expression (TPM > 1) in at least one tissue/stage within the chr 9:40 Mb-75 Mb TAD. Of the 370 genes, 248 (67%) are expressed during E8.5-10.5, and 15 are genes known to cause NTDs when mutated in mice, including *Bsb4*, *Foxb1*, *Aldh1a2*, *Nedd4* (Supplementary Fig. 2C). We leveraged the 43 stringent SNV hits as genetic markers in order to narrow down on the risk-variant. With additional exome sequencing we discovered 2727 different protein-affecting variants on chromosome 9 between the B6 reference genome and alternative 129S6 genome. Within the chr9:57 Mb-75 Mb locus there are 61 variants that are predicted to have a “moderate” to “high” impact on protein function (Supplementary Fig 2C).

Zooming down to the 0.9 Mb QTL peak (chr9:65,278,433-66,220,818) region, we found 1 “high” and 8 “moderate” impact variants in 9 genes, of which 5 are expressed and only one previously classified NTD candidate gene, *Snx1*, near the center (Fig. 2B and Supplementary Fig. 2C). *Snx1* encodes sorting nexin 1, a membrane protein involved in endocytosis and cellular trafficking (*42, 43*). We found two variants in *Snx1*; an inframe insertion encoding for a glutamic acid duplication (G137GG) in a stretch of already 4 other glutamic acids, and a G to A missense mutation resulting in a proline to serine variant (P117S). The *Snx1* variants are flanked by SNVs in the 5’ UTR and intron 2 of neighboring genes *Csnk1g1* and *Dapk2*, respectively (Fig. 2C). 5’UTRs contain regulatory sequences, and mutations or variants herein are still poorly understood, but can possibly affect transcription via interaction with the transcriptional machinery (*44*). Additionally, a publicly available dataset based on chromatin interaction analysis with paired-end tags mediated by RNA polymerase II (RNAP2 ChIA-PET), a technique to map cis-interactions between promoter regions, reported a functional interaction between the promoters of *Csnk1g1* and *Snx1* in mouse NPCs. This interaction coincides with CTCF binding sites, suggesting that both genes may be co-regulated via a chromatin loop (Fig. 2C). Both *Csnk1g1* and *Snx1* are bound by TET1 in both ESCs and EpiLCs, suggesting that they are direct TET1-regulated targets (Fig. 2C), and belong to gene families implicated in WNT signaling and secretion (*13, 45, 46*). Collectively, these observations suggest that either or both *Snx1* and *Csnk1g1* are candidate SNV-associated modifier genes, which in concert with *Tet1* loss can evoke a disruption in NTC.

**Fig. 2.**
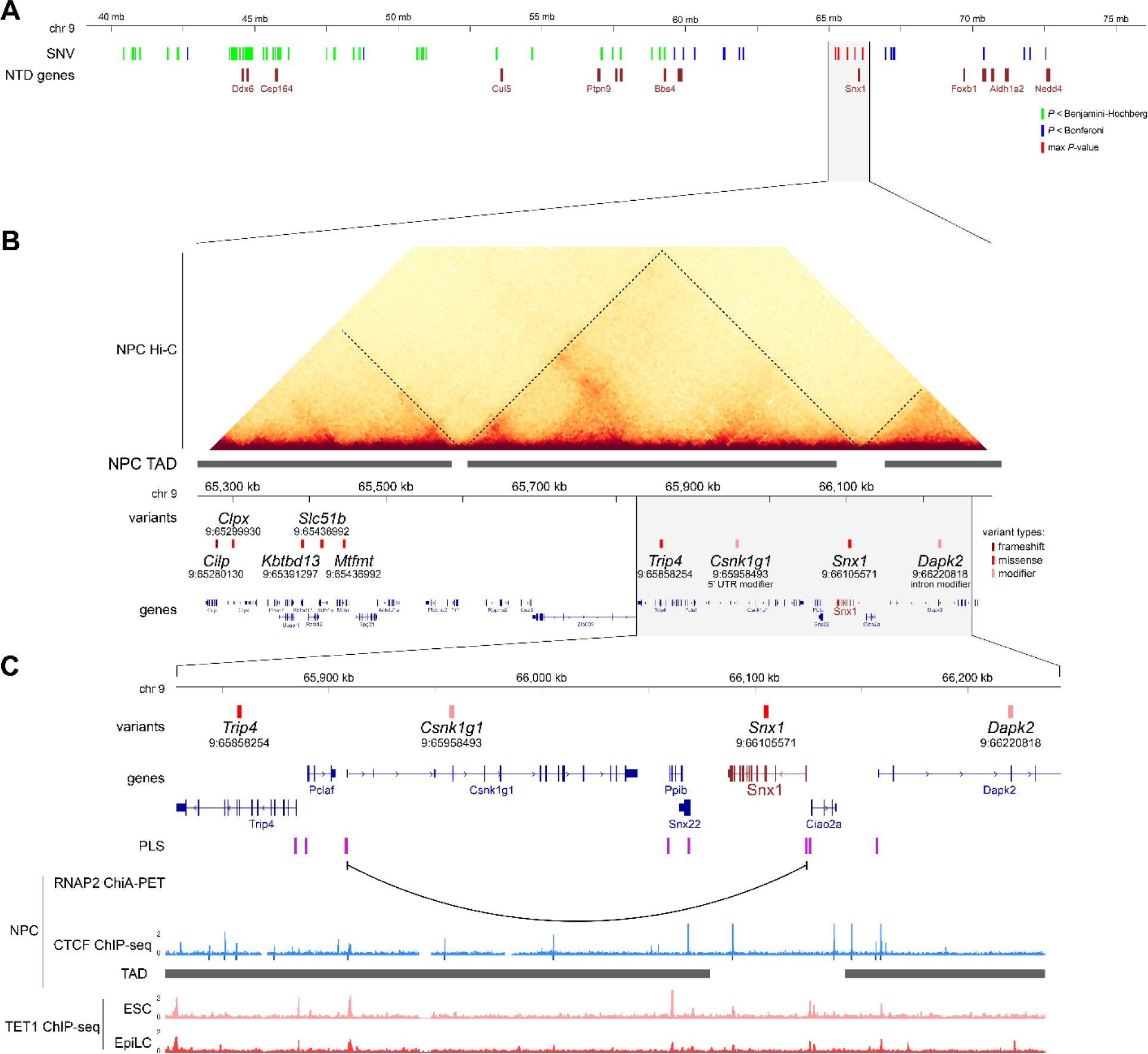
Candidate modifier genes at the QTL peak. (**A**), Genomic location of 216 SNVs above the Benjamini-Hochberg significance threshold within a 35Mb region on chromosome 9, relative to 15 NTD-related genes highlighted below in dark red. (**B**), Hi-C contact matrices (read densities in 5k bins) showing topology associated domains (marked by dashed lines) in mouse NPCs (*40*) over a 1 Mb region showing variants with a “high” or “moderate” impact on protein function. Gene annotations are shown below. (**C**), RNAP2 ChIA-PET interaction (*110*) and CTCF ChIP-seq in NPCs (dark blue bars below denote detected peaks of IDR p-value < 0.05), TADs (*40*), and TET1-ChIP-seq signals in ESCs (*35*) and EpiLCs (*33*) over the gene loci in the proximity of *Snx1* variant.

### Loss of *Tet1*^-/-^ in the embryonic lineage contributes cell non-autonomously to NTDs

NTC, a finely orchestrated process, is governed by a multitude of cellular signaling pathways, encompassing both canonical and non-canonical Wnt signaling (*5, 19, 47–49*). Our recent study revealed TET1’s role in regulating Wnt signaling, specifically through the regulation of a canonical Wnt repressor *Tcf7l1*, during the onset of germ layer lineage bifurcation (*34*). These experiments were performed using ESCs derived from the NTD-resistant B6129S6F1 background (*34, 35*). To interrogate whether strain differences influence TET1’s regulation of the cellular responsiveness to developmental signaling, potentially contributing to the etiology of NTDs, we derived another 6 independent male ESC lines (three *Tet1*^-/-^ and three *Tet1*^+/+^) from 129S6.B6-*Tet1*^tm1Koh^ blastocysts (*50*) to compare with the previous ESC F1 lines. These 129S6.B6 lines, juxtaposed with the previously established F1 ESC lines, were genotyped for the QTL presence to assess their strain-specific genetic makeup. As expected, the 129S6.B6 lines are homozygous for the 129S6 risk allele, while all F1 cells are heterozygous for both 129S6 and B6 alleles (Supplementary Fig. 3A).

To probe the potential of the new 129S6.B6 strain *Tet1* KO ESC lines in generating embryos with NTDs, we fluorescently labeled one 129S6.B6 *Tet1* KO ESC line (KO31) with lentiviral GFP and microinjected the cells as donors into tetraploid (4n) blastocysts (Supplementary Fig. 3B). In a tetraploid complementation assay, host cells from the tetraploid blastocyst are restricted in development to become only extra embryonic trophoblast-derived cells, so that the embryo is generated entirely from donor ESCs (Fig. 3A). Following the implantation of chimeric blastocysts into pseudo-pregnant mice, embryos were collected at E12.5. Although the efficiency of this method was low, producing only one embryo out of 13 deciduae recovered in an experiment, the embryo was GFP+ and clearly had an NTD (Fig. 3B). In contrast, NTDs were not observed in embryos derived from v6.5-*Tet1*^-/-^ ESC lines (50% B6; 50% 129S4) through tetraploid complementation, consistent with NTD resistance in F1 hybrid strains (*51*). This result confirms that the NTD phenotype results from *Tet1* loss-of-function in the embryonic lineage and is independent of its role in extra- embryonic development.

**Fig. 3.**
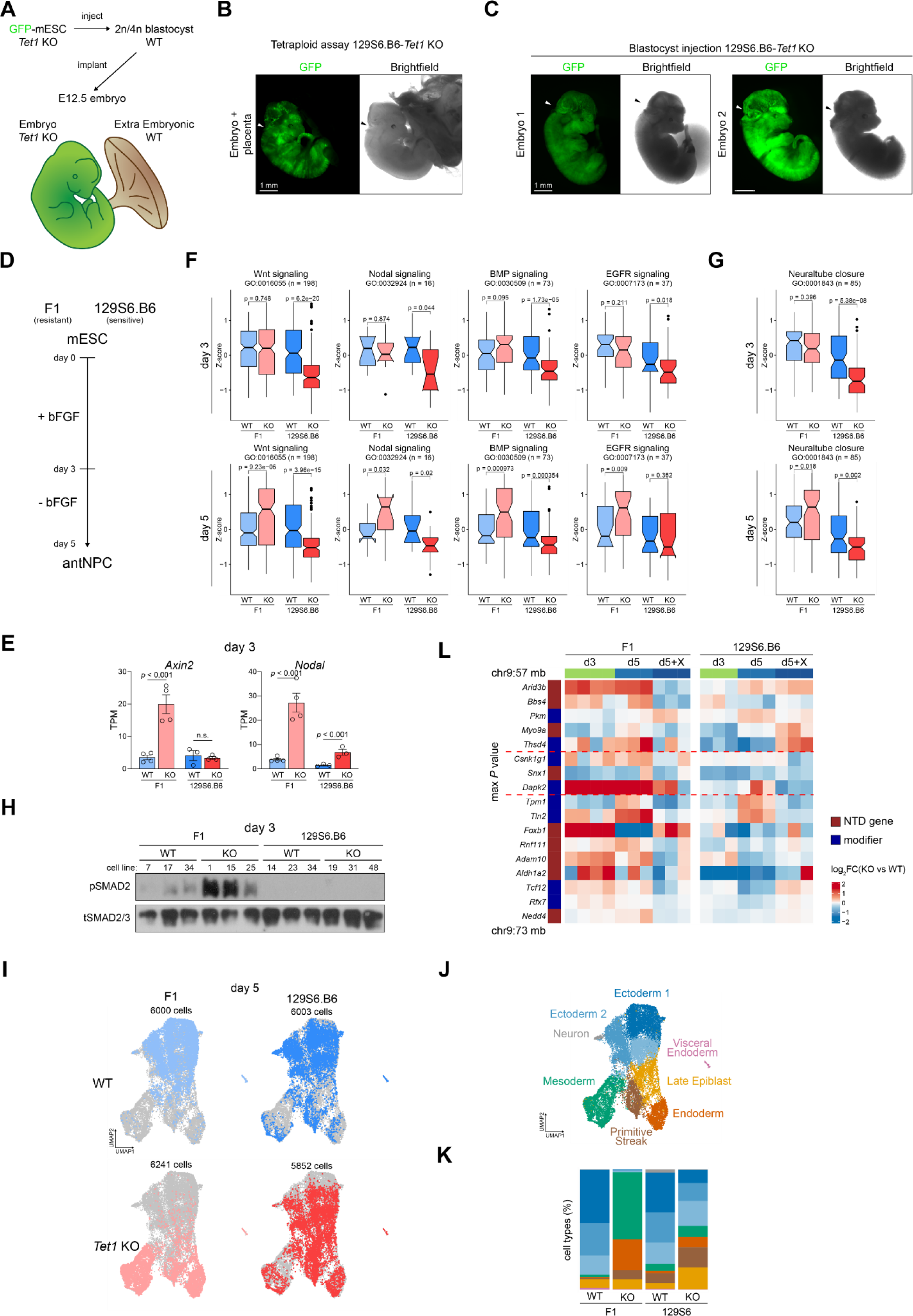
Strain differences in sensitivity to developmental signaling recapitulated by NTD- resistant and -sensitive strains of ESC lines. (**A**), Overview of experimental strategy to inject GFP+ *Tet1*^-/-^ mESCs into diploid or tetraploid host blastocysts to assess the contribution of donor cells in embryo-chimera. In a tetraploid blastocyst-chimera, the embryo proper is derived entirely from injected donor *Tet1*^-/-^ cells, while the host tetraploid cells are restricted to form the placenta. (**B**), Fluorescent (left) and brightfield (right) images of an embryo with placenta collected from a tetraploid chimera, showing GFP+ cells only in the embryo. Arrow indicates NTD. Scale bar is 1 mm. (**C**), Fluorescent and brightfield images of two chimeras with NTDs collected from diploid blastocyst injection. Arrow indicates exencephaly. Scale bar is 1 mm. (**D**), Schematic of *in vitro* differentiation to convert mouse ESCs into antNPCs (*34*). bFGF, basic fibroblast growth factor. (**E**), Gene expression of *Axin2* and *Nodal* measured in RNA-seq TPM on day 3 of differentiation. (**F**) and (**G**), Box- whisker plots of RNA-seq expression Z-scores of genes constituting each indicated GO- term in signaling (F) and NTC (G) in two strains of *Tet1*^-/-^ and *Tet1*^+/+^ cells, F1 (NTD- resistant) and 129S6.B6 (NTD-sensitive) background, collected on day 3 and day 5 of differentiation. Two independent differentiations per strain were performed using F1 *Tet1*^+/+^ and *Tet1*^-/-^ (*n*=4) and 129S6.B6 *Tet1*^+/+^ and *Tet1*^-/-^ (*n*=3) biological replicate ESC lines. Number of genes per GO term is indicated in parenthesis. The box represents the interquartile range (IQR), with the bottom and top edges corresponding to the first quartile (Q1, 25%) and the third quartile (Q3, 75%), respectively. The line inside the box represents the median (Q2, 50%) of the data. Whiskers extend from the box to a maximum of 1.5 times the IQR, and individual data points beyond the whiskers are considered outliers. (**H**), Western blot of phospho (p)SMAD2 and total (t)SMAD2/3 on day 3 of differentiation. Numbers above each lane denote identities of independent blastocyst-derived cell line per genotype and strain. *Csnk1g1*, *Snx1* and *Dapk2* are genes with the max *P* value and highlighted with red dashed lines. (**I**), UMAP plot for scRNA-seq of *Tet1*^+/+^ and *Tet1*^-/-^ F1 and 129S6.B6 cells on day 5 of differentiation. (**J**), UMAP clusters annotated for cell lineage. (**K**), Proportion of each cell type per sample. (**L**), RNA-seq normalized expression heatmap of genes, including 9 NTD-implicated genes, found within a 15-Mb region surrounding the QTL peak and exhibiting expression (TPM>1) in at least one experimental group during *in vitro* differentiation of 2 strains of WT and *Tet1* KO ESCs to NPCs. The log2FC is calculated based on transcripts per million (TPM) of KO vs WT.

To increase the efficiency of embryo-chimera production, we subsequently performed injections of 129S6.B6-*Tet1* KO ESCs into diploid blastocysts, generating chimeric embryos with a mosaic of host *Tet1*^+/+^ and donor *Tet1*^-/-^ cells. Of 20 deciduae recovered, 13 contained embryos of which 9 were chimeric with varying degrees of GFP^+^ contribution. Remarkedly, two of the 9 chimeras exhibited exencephaly (Fig. 3C), suggesting that the presence of *Tet1*^-/-^ donor cells among *Tet1*^+/+^ recipient cells disrupts NTC in a cell non-autonomous manner. Altogether, these results affirm that the distinct NTD susceptibilities induced by TET1 dysfunction in F1 versus 129S6.B6 backgrounds can be faithfully recapitulated in the corresponding ESC lines. These lines stand as potent *in vitro* models that provide novel insights into the underlying mechanisms governing NTC and strain-dependent susceptibility to NTDs.

### NTD-sensitive 129S6.B6 ESCs exhibit impaired developmental signaling response during differentiation

To discern disparities in differentiation potential between the NTD-resistant F1 and NTD-sensitive 129S6.B6 ESCs, we used a serum-free non-directed differentiation protocol to guide ESCs “default” conversion into anterior NPCs (antNPCs) in the absence of exogenous morphogen or signaling inhibitors (*34, 35, 52, 53*). In our previous studies with F1 cells, we observed that *Tet1*^-/-^ cells switched in lineage towards mesoderm and endoderm as a result of ectopic activation of canonical Wnt and Nodal signaling, whereas *Tet1*^+/+^ cells form exclusively definitive ectoderm (*34*).

We performed bulk mRNA-seq with both 129S6.B6-*Tet1*^-/-^ and *Tet1*^+/+^ cells on day 3 and day 5 of differentiation, corresponding to gastrulation onset and post-gastrulation stages respectively, and compared with previous mRNA-seq analysis of the F1 cells (Fig. 3D). Principal component analysis (PCA) showed that 129S6.B6-*Tet1*^-/-^ cells differentiate into neuroectodermal cells akin to *Tet1*^+/+^ counterparts but distinct from F1-*Tet1*^-/-^ cells (Supplementary Fig. 3C). In both backgrounds, we detected roughly equal numbers of upregulated and downregulated differentially expressed genes (DEGs) between *Tet1*^-/-^ vs *Tet1*^+/+^ (using FDR adjusted p-value < 0.05) on both day 3 and day 5, although there were more DEGs in the F1 (total DEGs d3 = 2327, d5 = 3281) than in the 129S6.B6 (total DEGs d3 = 1448, d5 = 1826) pairwise comparisons (Supplementary Fig. 3D). In the 129S6.B6 background, DEGs upregulated on day 3 were associated with general cellular processes, including cell adhesion, but clearly lacked enrichment for GO terms associated with primitive streak formation and Wnt-signaling observed in the F1 strain. Down-regulated DEGs in both F1 and 129S6 strains on day 5 enriched for neurogenesis associated terms, including forebrain development, but the term “tube formation” was additionally enriched in 129S6 cells (Supplementary Data S1). Notably, Wnt (*Axin2*, *Sp5*, *Lrp4*, *Wnt9b*, *Tcf7*) and Nodal (*Nodal*, *Lefty1*, *Tdgf1*) targets were up-regulated in F1- *Tet1*^-/-^ cells, but not in 129S6.B6-*Tet1*^-/-^ cells (Fig. 3E, Supplementary Fig. 3E).

Comparing the expression level of all genes annotated in the GO terms for various signaling pathways revealed a significant reduction in expression in 129S6.B6-*Tet1*^-/-^ cells compared to *Tet1*^+/+^ cells, particularly in Wnt, Nodal, BMP and EGFP signaling on day 3 (Fig. 3F, Supplementary Fig. 3F-G). The term “neural tube closure” was also significantly downregulated on day 3 in 129S6.B6 *Tet1*^-/-^ cells, but not in F1 (Fig. 3G). This trend persisted until day 5, when the F1-*Tet1*^-/-^ cells upregulated all pathways significantly, while 129S6-*Tet1*^-/-^ cells down-regulated all pathways instead (Fig. 3F, Supplementary Fig. 3F,G). Genes associated with canonical- and non-canonical Wnt signaling pathways were consistently downregulated in 129S6.B6-*Tet1*^-/-^ cells throughout differentiation (Supplementary Fig. 3H). This was further validated through western blots for phosphorylated (p)SMAD2. On day 3, an increase in Smad2 phosphorylation was observed in F1- *Tet1*^-/-^ cells, while not in the 129S6.B6 cells (Fig. 3H).

To validate the lineage fate of 129S6.B6-*Tet1*^+/+^ and *Tet1*^-/-^ cells upon non-directed differentiation, we performed scRNA-seq on day 5 in the absence of Wnt inhibition. In contrast to our previous scRNA-seq of F1 cells, over 50% of 129S6.B6-*Tet1*^-/-^ cells formed ectoderm clusters, with only around 20% diverging into mesoderm and endoderm by day 5 (Fig. 3I-K, Supplementary Fig. 3I,J). While both strains displayed a similar extent of *Tcf7l1* expression regulated by TET1, in the 129S6.B6 strain, an impaired capacity to activate signaling pathway genes leads to a default route of neural induction independent of TET1. As *Tet1*^+/+^ cells on day 5 resemble the E8.5 neuroepithelium post-gastrulation, coinciding with the initiation of neural tube closure, these data suggest that impaired Wnt and Nodal signaling in *Tet1* KO cells of the 129S6.B6 strain may subsequently result in NTDs.

Using these strain-comparative RNA-seq datasets, we also interrogated expression patterns of genes within the chr 9 QTL. Focusing on the chr9:57Mb-75Mb region flanked by *Pkm* and *Rfx7* SNVs, which contiguously surpassed the stringent Bonferroni threshold (Fig. 1I,J), we charted the time-course expression of 17 genes associated with modifier SNVs, of which 9 are implicated in NTDs, in both strains. Interestingly, we observed a co-regulated pattern of gene activation during differentiation, highly sensitive to WNT inhibition by XAV939, in the NTD-resistant F1 strain (Fig. 3L). In contrast, gene activation of the entire cluster was severely impaired in the NTD-sensitive 129S6.B6 strain. A quantitative PCR measurement of *Csnk1g1*, *Snx1* and *Dapk2* relative gene expression selectively in additional experimental replicates validated *Dapk2* to be significantly up- regulated in *Tet1* KO NPCs in the F1, but not in the 129S6.B6 strain (Supp Fig. 3K). These results implicate the chr9 QTL locus as a hot-spot of co-regulated Wnt-sensitive genes in a hybrid strain that is resistant to NTDs, but which are collectively impaired in their sensitivity to Wnt signaling in the 129S6.B6 strain to result in increased susceptibility to NTDs.

### NTDs in *Tet1*^-/-^ embryos are resistant to FA supplementation across strains

Given that FA is a major protection against NTD risk, we asked whether the penetrance of NTDs caused by loss of *Tet1* is sensitive to maternal folate status across the different mouse strains we have created. We initially designed experiments to explore the potential of surplus FA to mitigate NTD penetrance in the highly susceptible CD1-*Tet1^tm1Koh^* KO mice, and conversely, whether folate deprivation could elevate NTD rates in less sensitive congenic B6-*Tet1 ^tm1Koh^* KO strain (Supplementary Fig. 4A). The CD1-*Tet1*^+/-^ dams underwent a diet adaptation to 2.1 ppm FA for 4 weeks, reduced from 7 ppm in vitamin-enriched chow regularly used in our mouse facility, prior to timed-mating with CD1-*Tet1*^+/-^ stud. Following a previously published study, 10 mg/kg FA (or PBS as vehicle control) were then intraperitoneally injected daily from E5.5, with embryos collected at E11.5 (*54*) (Supplementary Fig. 4A). Direct FA injection did not result in lower NTD rates in CD1- *Tet1*^-/-^ embryos compared to controls and might even have slightly exacerbated NTD penetrance, although the difference did not reach statistical significance (*p* = 0.22, n = 56 from 20 litters in FA group, n = 54 from 19 litters in PBS group) (Supplementary Fig. 4B). To address the possibility that the effects of folate cycling may be uncoupled with that of methionine, we also tested injections of methionine at 70 mg/kg in CD1-*Tet1*^+/-^ dams, but this showed no effect on NTD rates in KO offspring (Supplementary Fig. 4A-B). To perform FA deprivation in B6 mice, we adapted B6-*Tet1*^+/-^ dams to a diet containing 0.1 ppm FA for 4 weeks before mating with *Tet1*^+/-^ studs. To further eliminate folate synthesis by intestinal flora, the 0.1 ppm FA diet was supplemented with an antibiotic, 1% succinyl sulfathiazole (SST), as used in previous studies (*54*). The control diet for this set of experiments contained 2.7 ppm FA plus 1% SST. Here as well, FA deficiency did not increase NTD rates in B6- *Tet1*^-/-^ embryos (*p* = 0.5, n = 32 from 19 litters in 0.1 ppm FA group, n = 32 from 16 litters in 2.7 ppm FA group.) (Supplementary Fig. 4C). These observations led us to conclude that the NTDs caused by loss of TET1 are non-responsive to FA dietary changes across strains.

Folate 1C cycling provides substrates for methylation reactions, but TET dioxygenases erase DNA methylation to counteract excessive DNMT activities; we therefore reasoned that loss of TET1 activity may impact 1C metabolism in embryos. To gain a comprehensive view of steady-state metabolic changes induced in *Tet1* WT and KO embryos in response to maternal FA status, we performed an untargeted high-performance liquid chromatography with tandem mass spectrometry (HPLC-MS/MS) analysis of 598 metabolites in the E11.5 whole embryos collected. Notably, FA injection in CD1-*Tet1*^+/+^ embryos resulted in a significant increase in FA and a global increase in metabolites belonging to phospholipid and carnitine pathways, a response not mirrored in *Tet1*^-/-^ embryos (Supplementary Fig. 4D). Conversely, FA deprivation in B6-*Tet1*^+/+^ embryos resulted in down-regulation of phospholipids, but an increase in carnitine metabolites; while *Tet1*^-/-^ embryos remained nonresponsive (Supplementary Fig. 4E). These differences in metabolomic response between WT and *Tet1* KO embryos were observed independently of embryo sex and NTD presence, suggesting that the loss of *Tet1* is the dominant factor disrupting the homeostatic regulation of 1C metabolism in response to nutritional insults post-neurulation. While either excess or deficient FA status will drive up carnitine metabolism in WT (but not in *Tet1* KO) embryos, levels of phospholipid metabolites appear positively correlated with FA status in a manner dependent on *Tet1*, in line with folate being a source of one-carbon for lipid metabolism (da Silva et al. 2014).

### Maternal folate status interacts with *Tet1* gene dosage to induce congenital brain malformations

To further probe how adverse maternal dietary folate status interacts with *Tet1* dysfunction to affect offspring development, we subsequently investigated the impact of excess and depleted maternal FA on E11.5 *Tet1*^tm1Koh^ embryos within the 129S6 congenic (129S6.Cg) strain of mice after 13 generations of backcrossing to 129S6, to eliminate the confounder of genetic heterogeneity. Based on the higher NTD penetrance observed in 129S6.B6 incipient congenic *Tet1* KO embryos, we also reason that the 129S6 background would contain the bulk of genetic variants that increase susceptibilities for developing embryonic defects compared to B6. The low NTD penetrance in 129S6.Cg-*Tet1*^-/-^ mice also prompted us to explore structural defects other than NTDs at E11.5 post-neurulation. According to “US FDA 2005 Guideline for Industry” to convert human dose to animal dose, an adult mouse consuming 3-5 g chow daily on a 3 ppm FA diet has an intake “equivalent” to 0.11-0.18 mg of FA per day in human (US CDC recommended dietary allowance is 0.4 mg) (*23*). On this basis, a high FA diet containing 30 ppm FA was designed to achieve a 10-fold FA excess in rodents comparable to >1 mg daily intake in human. To establish an accurate/known level of FA availability from the diet and to reduce experimental variability , all custom diets were supplemented with 1% SST to eliminate folate contribution from gut microbiota, as described above (Fig. 4A) and in line with previous studies (Burren et al. 2008; Christensen et al. 2015; Cosin-Tomas et al. 2020), , because the microbiota is a major source of folate in rodents. Using CD1-*Tet1^tm1Koh^* mice, we verified that NTD rates of *Tet1*^-/-^ (KO) were not significantly altered by custom diets containing 3 or 7 ppm FA (matching levels in regular chow) with or without SST (Supplementary Fig. 4F).

**Fig. 4.**
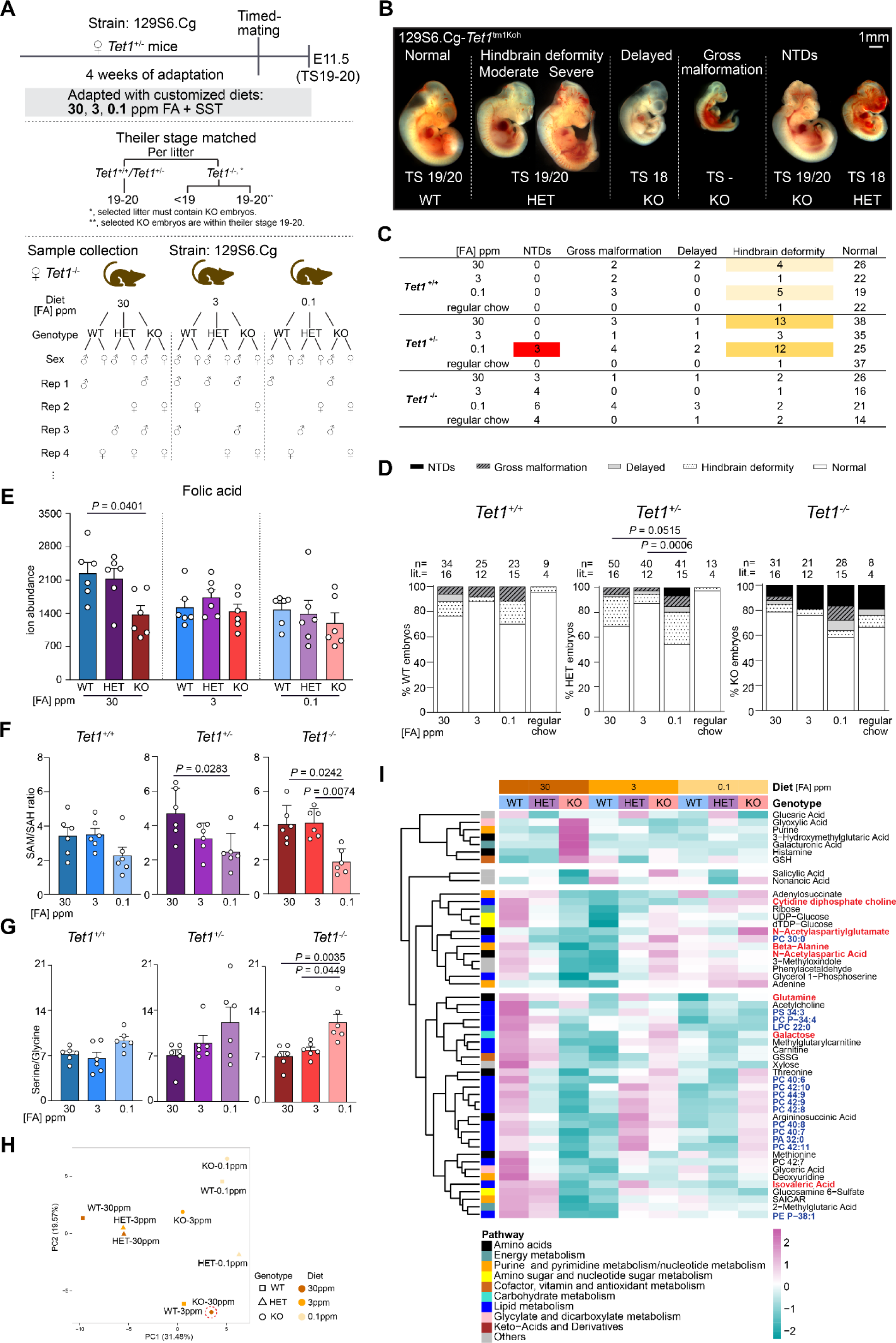
Phenotypic and metabolomic changes in 129S6.Cg embryos in response to maternal dietary FA excess or depletion, interacting with the loss of Tet1. (**A**), Schematic of protocol adapting *Tet1*^+/-^ dams to custom diet containing surplus (30 ppm), normal (3 ppm, control) or depleted (0.1 ppm) FA (top), and sample selection criteria to match KO with littermate WT and/or HET embryos by Theiler stage (TS19-20) and sex per biological replicate (middle and bottom). All three diets contain SST, succinyl sulfathiazole, an antibiotic to eliminate folate contribution from gut microbiota. For each downstream assay, we collect 3-6 biological replicates from independent litters, comprising similar numbers of male and female replicates. (**B**), Representative images of E11.5 embryos of the 129S6.B6- *Tet1*^tm1Koh^ strain, illustrating distinct phenotypes observed as a result of maternal dietary FA modulation. Theiler stage and *Tet1* genotype are indicated below each embryo. Scale bar is 10 mm. (**C**), Tabulation of embryo counts by phenotypes observed in WT, HET and KO embryos of the 129S6.B6-*Tet1*^tm1Koh^ strain, exposed to FA-modified maternal diet. The regular chow contains 7 ppm FA without SST. (**D**), Phenotypic rates observed in WT, HET and KO embryos of the 129S6.B6-*Tet1*^tm1Koh^ strain exposed to FA-modified maternal diet. lit. = litter counts, n = embryo counts. *P* values are calculated by Fisher’s Exact test. (**E**-**G**), HPLC-MS/MS detection of unmetabolized FA (E), ratio of SAM/SAH (F), and serine/glycine (G) in whole embryos. Data are mean ± SEM from n=6 individual embryos per group, consisting of 3 males and 3 females, except WT in 30 ppm FA and KO in 3 ppm FA groups which were made up of 4 females and 2 males. * *P* < 0.05 by ANOVA. (**H**), Principal Component Analysis (PCA) of grouped WT, HET and KO samples exposed to the three FA diets. (**I**), Heatmap of metabolites with significant changes in abundance as a result of interactions between folate status and *Tet1* genotype (2-way ANOVA interaction *P* < 0.05). Dendrogram shows k-means clustering and scaled counts are Z-score of ion abundance values. Two samples were extreme outliers within their groups in the PCA and excluded from differential analysis: one HET female embryo in 3 ppm FA (n=5) and one WT male embryo in 30 ppm FA, resulting in n=5 in these two groups. Phospholipids are highlighted with blue in bold, neurotransmitters and neuronal endocytosis/exocytosis are highlighted with red in bold, respectively.

Custom diets containing 30 ppm (excess), 3 ppm (regular) or 0.1 ppm (depleted) FA (with 1% SST) were administered to timed-pregnant 129S6.Cg-*Tet1*^+/-^ dams, and embryos were phenotyped at E11.5 (Fig. 4A). An additional reference group was kept on standard chow enriched with 7 ppm FA (without antibiotics). The use of custom diet introduced a basal rate of gross malformation 5-8%) and increased resorption rates especially under FA deprivation (Fig. 4B-D and Supplementary Fig. 5A). We regarded these as a side-effect of antibiotic supplementation in the custom diets not attributed to FA status. Nonetheless, we observed an increased incidence of morphological abnormalities (tissue distortion) at the hindbrain-midbrain junctures (‘hindbrain deformity”) in *Tet1*^+/+^ (WT) and *Tet1*^+/-^ (HET) embryos exposed to FA excess or deprivation (Fig. 4B-D). These observations would be consistent with neurodevelopmental deficits reported in rodent offspring exposed to adverse (excess and deficient) maternal folate status (*55, 56*). The penetrance of such “hindbrain deformities” were notably exacerbated among *Tet1*^+/-^ embryos when exposed to excess and depleted FA (20-30%, compared to 10-15% among *Tet1*^+/+^ embryos), and significantly above the basal incidences among *Tet1*^+/-^ (< 5%) in the control group (3 ppm FA +SST) and in regular chow (7 ppm FA). The phenotypes were superseded by open NTDs in *Tet1*^-/-^ (KO) embryos, observed at similar rates under all dietary conditions, suggesting that the hindbrain deformities may be precursors of NTDs triggered by *Tet1* insufficiency (Fig. 4C-D).

Strikingly, we found 3 NTD cases in two litters among 41 *Tet1*^+/-^ (HET) embryos examined (7%), specifically in the FA depleted groups. Previously, NTDs were never observed in *Tet1*^+/-^ embryos under standard chow conditions (in over 1000 experimental time-pregnancies conducted using *Tet1*^tm1Koh^ strains), suggesting that folate depletion coupled with *Tet1* haploinsufficiency can increase the susceptibility to NTDs. A re-examination of previous experiments in B6 and CD1 mice treated with FA-modified diets revealed subtle malformations not seen in regular chow experiments. In B6-*Tet1*^+/-^ embryos in the FA-deprived cohort, a small subset displayed missing pigmentation and malformation of the eyes, resembling anophthalmia and microphthalmia, at a rate of 11%, above that (6%) among WT embryos, including one NTD case (Supplementary Fig. 4C, G). In CD1 mice, hindbrain deformity phenotypes were observed at low rates of less than 3% in HET and ∼5% in KO embryos, but not at all in WT, when exposed to excess and deprived FA (Supplementary Fig. 4G, H). These observations illuminate that under nutritional stress, such as folate deprivation, *Tet1* haploinsufficiency can heighten the risks for congenital malformations including NTDs.

### Adverse maternal folate status under *Tet1* deficiency affects phospholipid metabolism

To understand how maternal FA intake and *Tet1* gene dosage interact to shape cellular physiology, we proceeded to dissect molecular changes at the level of the metabolome, DNA methylome, and transcriptome. To profile the metabolomic changes, we again performed untargeted HPLC-MS/MS of 807 metabolites in whole E11.5 129S6.Cg-*Tet1*^tm1^ embryos per custom diet and *Tet1* genotype (3 x 3 experimental groups). In each biological replicate (n=6), *Tet1*^-/-^ embryos were stage- and sex-matched to littermate *Tet1*^+/+^ and/or *Tet1*^+/-^ embryos, excluding embryos with gross malformation and developmental delays (Fig. 4A). In our sample sets of 54 embryos (Supplementary Table S2), we intentionally included a few WT and HET embryos with mild deformities (but stage-matched based on somite counts) and selected against KO embryos with severe malformation. Detection of unmetabolized FA revealed a two-fold increase in *Tet1*^+/+^ and *Tet1*^+/-^ embryos under excess FA, but this increase was nullified in *Tet1*^-/-^ embryos (Fig. 4E). Conversely, FA-depletion in *Tet1*^+/-^ and *Tet1*^- /-^ embryos resulted in a reduced SAM/SAH ratio, a measure of the cellular methylation index, and an increased serine/glycine ratio, indicative of reduced rates in cycling of active folate species (Fig. 4F-G). These findings indicate a sensitivity of folate/methionine cycling towards *Tet1* gene dosage under conditions of folate deficiency.

Principal Component Analysis of 84 metabolites exhibiting significant changes in ion abundances in at least one experimental group by one-way ANOVA showed sample groups clustering by dietary FA status along the first principal component (PC1). As an exception, KO embryos exposed to excess FA (group 30 ppm FA_KO) separated from HET and WT embryos along both PC1 and PC2 (Fig. 4H). PCA of the differential metabolome in individual embryos showed two (TP12.3_3ppm_HET_female and TP25.6_30ppm_WT_male) to be extreme outliers from other replicates in their respective groups (Supplementary Fig. 5B); these two samples were excluded in the downstream analysis but otherwise did not alter the outcome. To further identify metabolites showing significant changes in abundance as a result of interactions with both diet and genotype, we used two-way analysis of variance (ANOVA). Interestingly, the analysis revealed that a majority (40.4%) of significantly changed metabolites belonged to pathways of lipid metabolism (Fig. 4i, Supplementary Fig. 5C). Specifically, several clusters including phosphatidylcholines (PCs), neurotransmitters, and metabolites involved in neuronal endocytosis/exocytosis, were significantly increased in *Tet1*^+/+^ and *Tet1*^+/-^ embryos under excess FA, but not in *Tet1*^-/-^ (Fig. 4I). Conversely, these PC compounds were reduced in *Tet1*^+/+^ and *Tet1*^+/-^ embryos relative to *Tet1*^-/-^ under FA depletion, strongly supporting that their cellular abundance are highly sensitive to the loss of TET1. The differences were observed independently of the embryo’s sex and phenotype and recapitulated the compromised metabolomic response of *Tet1*^-/-^ embryos to adverse folate status in the CD1 and B6 backgrounds (Supplementary Fig. 4D, Supplementary Fig. 5C). These findings underscore the intricate functional interactions between TET1 and folate one-carbon metabolism and suggest that FA-resistant NTDs observed in *Tet1* null embryos may stem from an impaired ability to metabolize or uptake FA.

### FA-sensitive differentially methylated regions caused by the loss of TET1 are enriched for neurodevelopmental loci

Given that adverse folate status can affect cellular methylation potential, we considered genome-wide DNA methylation levels as an epigenetic readout to discern the consequences of *Tet1* loss and dietary variations in FA. For this genomic analysis, we dissected E11.5 brain tissues from the most rostral aspect of the forebrain to the caudal aspect of the hindbrain above the otic vesicle, following ENCODE guidelines (Supplementary Fig. 6A), from 129S6.Cg mice exposed to all three FA diets. We selected individual embryos matched by somite counts, pairing a KO with one WT and/or HET of the same sex within a litter as a replicate unit (Fig. 4A). Our final sample set included a total of 36 embryos collected from 5 litters in each diet group, consisting of equivalent numbers of male and female replicates and representative of the phenotypes observed (Supplementary Table S2). To profile DNA methylation changes in this sample cohort cost-effectively, we performed enhanced reduced representation bisulfite sequencing (RRBS), which is based on enrichment of MspI and Taq1 restriction sites (*57*). From sequencing 20 million PE150 reads per sample, we obtained a mean sequencing depth of 8.7x covering about 2 million CpGs. The PCA projection and cluster analysis showed samples separating into two clusters along PC1 according to KO versus WT/HET genotype, indicating that complete loss of TET1 function, rather than diet, was the dominant driver of methylation differences (Supplementary Fig. 6B,C). One distinct outlier sample (HET embryo FA-depleted), coincidentally also the sample with the lowest coverage of 4.9x, was excluded in downstream analysis (Supplementary Fig. 6B).

Aggregate methylation level profiles across the gene structure of 21231 protein-coding genes, when stratified by *Tet1* genotypes, revealed a subtle decrease in methylation, particularly in 5-kb upstream and downstream flanking regions, exclusively in WT samples under FA depletion (Fig. 5A). To identify differentially methylated regions (DMRs) dependent on *Tet1* genotypes, we conducted pairwise comparison analysis between *Tet1*^-/-^ or *Tet1*^+/-^ vs *Tet1*^+/+^, per diet group. As expected, a majority (99.3%) of 33851 DMRs exhibited gains in DNA methylation (hyper DMRs) in KO versus WT across all diets (Fig. 5B). However, excess and depleted FA diets resulted in fewer hyper DMRs compared to the control diet, suggesting that adverse maternal folate levels impede DNA hypermethylation linked to TET1 deficiency. Similarly, in the comparison between HET and WT in normal FA diet, we also detected 6030 hyperDMRs, indicative of a *Tet1* gene dosage effect on DNA methylation, which were completely obliterated by modified FA levels. When we compared 30 ppm or 0.1 ppm with 3 ppm condition, stratified per genotype, to assess the dietary impact, we discovered >10-fold fewer DMRs, the majority (93.6% of a total of 7057) afflicting WT embryos under both excess and depleted FA diet (Fig. 5C). Excess FA predominantly induced hyperDMRs (65.3%) in WT embryos, consistent with the expected positive influence of folate cycling on DNA methylation pathways. The 857 hyperDMRs identified enriched for GO terms related to nervous system development and function, including “regulation of membrane potential”, “regulation of action potential” and “action potential” (Supplement Fig. 5D). In contrast, depleted FA induced gain and loss in methylation equally, with DMRs outnumbering those caused by excessive FA, suggesting that FA deprivation has a more extensive impact on the epigenome than FA excess. While hyperDMRs induced by FA deficiency in WT embryos associated with general developmental terms, the hypoDMRs enriched for Wnt signaling pathway (Supplement Fig. 5E).

**Fig. 5.**
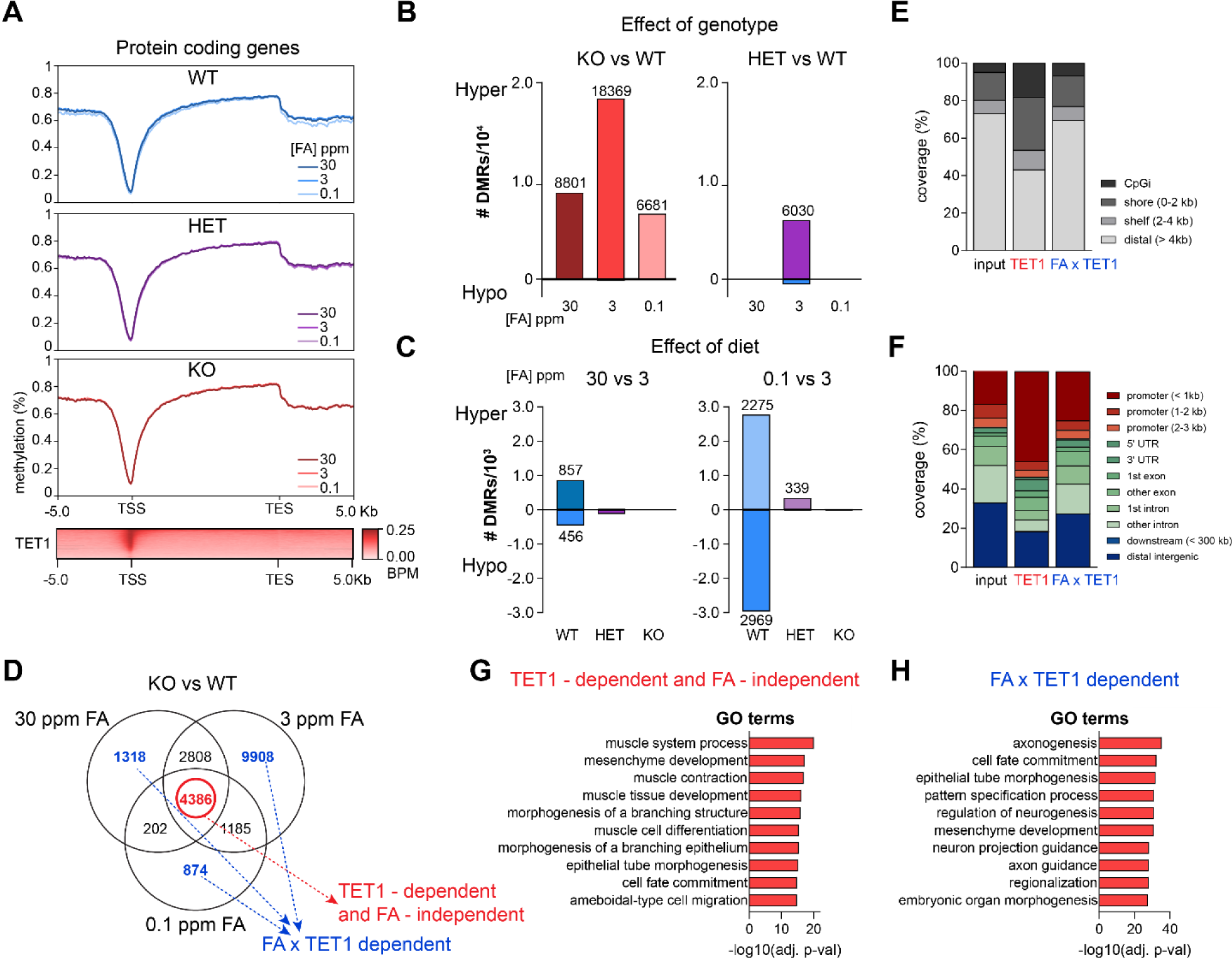
Genomic features of FA-sensitive differentially methylated regions interacting with the loss of TET1. (**A**), Profiles of aggregate methylation levels across the gene structure of 21231 protein coding genes in 30, 3 and 0.1 ppm FA stratified by *Tet1* genotypes. n=5, KO per diet group, matched to n=3 WT and n=6 HET under 30ppm FA, n=4 WT and n=3 HET under 3ppm FA, and n=2 WT and n=3 HET under 0.1ppm FA. Bottom, ChIP-seq profile of TET1 binding at the transcription start sites (TSS) of these genes. (**B**) and (**C**), Number of hyper and hypo differential methylated regions (DMRs) defined by *Tet1* genotype KO or HET vs WT pairwise comparisons (B) or by FA diet 30 ppm or 0.1 ppm vs 3 ppm pairwise comparisons (C) of E11.5 embryonic brain samples. (**D**), Venn diagram of hyper DMRs defined by KO vs WT pairwise comparisons of E11.5 embryonic brain samples per diet in (B). DMRs common to all diets are circled in with red and classified as TET1 specific DMRs, whereas the collective set of DMRs unique to each diet are colored in blue and classified as FA x TET1 DMRs. (**E**) and (**F**), Distribution by CpG island proximity (E) and gene feature annotation (F) of TET1-specific and FA x TET1 DMRs. (**G**) and (**H**), GO analysis of TET1- specific (G), and FA x TET1 (H) DMRs.

Subsequently, we focused on hyperDMRs resulting from *Tet1* KO across the three custom FA diets, distinguishing those regions resulting solely from loss of *Tet1* from those sensitive to modulation by FA status. By comparing the three hyper DMRs subsets, we identified 4386 regions exhibiting hypermethylation consistently in all three custom FA diets, indicating that these DMRs resulted exclusively from loss of TET1 (i.e. TET1-dependent and FA-independent), unaffected by diet variations (Fig. 5D). In contrast, hyperDMRs exclusive to each dietary group represented regions regulated by FA and TET1 interactions (i.e. FA x TET1 dependent). When annotated by genomic features, TET1-dependent DMRs were notably enriched in the CpG islands and CpG shores, closely associated with gene-proximal promoter regions while FA x TET1 DMRs showed enrichment on the promoter regions to a lesser extent (Fig. 5E-F). Interestingly, GO analysis of TET1-dependent DMRs revealed an enrichment in mesoderm-related terms, including “muscle system process”, “mesenchyme development” and “muscle contraction” (Fig. 5G). In contrast, FA x TET1 DMRs enriched for ectoderm-related terms, including “axonogenesis“, “cell fate commitment,” and “epithelial tube morphogenesis” (Fig. 5H). Collectively, these findings unravel an interesting interaction between adverse FA status and TET1 dysfunction that induces aberrant DNA methylation patterns that converge at genomic loci enriched for neurodevelopmental functions.

### Excess maternal folate interaction with loss of TET1 down-regulates membrane transporter expression

To bridge the observed metabolomic and DNA methylation shifts described above with gene expression, we performed bulk RNA-seq on E11.5 brains collected from 129S6.Cg mice exposed to the three custom FA diets. Again, 4-6 biological replicates (comprising male and female littermate pairings) were collected by sex- and stage-matching KO embryos to littermate WT and/or HET embryos, to compile a total of 40 samples for analysis (Supplementary Data S1). PCA projection showed that all samples clustered close together independently of diet group and genotype, except two samples (one *Tet1*^+/+^and one *Tet1*^-/-^ embryo in excess FA groups) which were from malformed embryos and henceforth excluded in downstream analysis (Supplementary Fig. 7A). To find differentially expressed genes (DEGs) dependent on *Tet1* genotypes, we first performed pairwise comparisons between *Tet1*^-/-^ vs *Tet1*^+/+^, or *Tet1*^+/-^ vs *Tet1*^+/+^, within each diet group. Interestingly, more than 79% of DEGs (FDR adjusted *p*-value < 0.05) were in the excess-FA group comparing *Tet1*^-/-^ vs *Tet1*^+/+^ and almost all downregulated (176 down, 17 up) in *Tet1*^-/-^ (Fig. 6A). The downregulated DEGs were enriched in GO terms related to neurotransmission, including “vesicle-mediated transport in synapse”, “neurotransmitter secretion”, and “protein localization to synapse” (Fig. 6B). Many of these affected genes encode solute carriers (SLC) facilitating transmembrane transport of L-glutamate and γ-amniobutyric acid (GABA), both crucial neurotransmitters in emotion and cognition (Fig. 6C). Pairwise comparisons between *Tet1*^+/-^ and *Tet1*^+/+^ embryos per diet group yielded minimal DEGs (only 19 and 32 DEGs in either group), indicating *Tet1* haploinsufficiency has negligible impacts on gene expression in embryonic brains under unaltered dietary conditions (Fig. 6A).

**Fig. 6.**
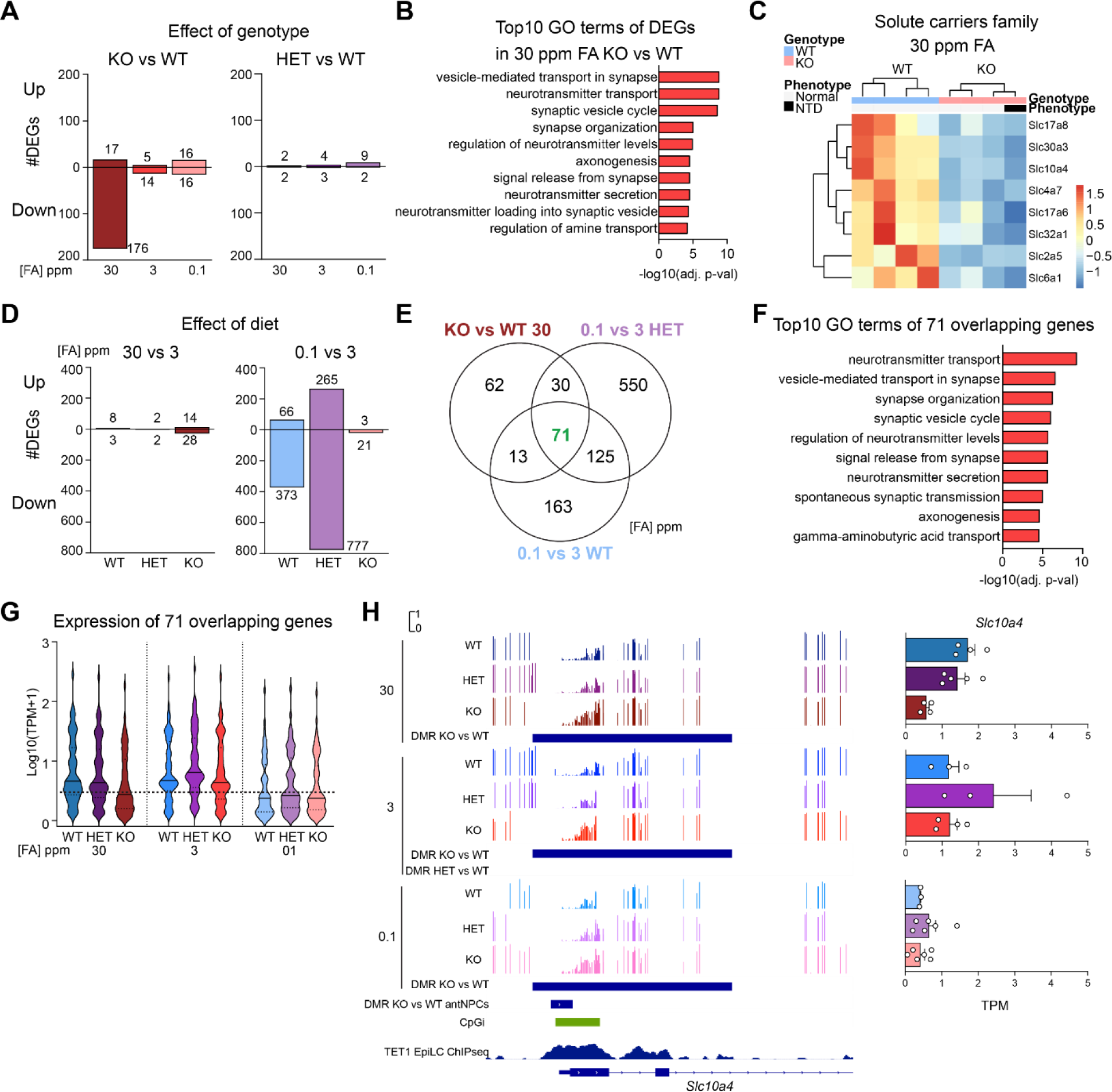
Down-regulation of membrane transporter expression by an interaction of adverse maternal folate interaction with loss of TET1. (**A**), Number of up and down-regulated RNA-seq differential expressed genes (DEGs) defined by genotype *Tet1* KO or HET vs WT pairwise comparisons of E11.5 embryonic brain samples per diet. Differential cut-off was FDR-adjusted *P* value < 0.05. (**B**), GO analysis of the 176 down-regulated DEGs defined in KO vs WT exposed to 30 ppm FA, as shown in (A). (**C**), Heatmap of RNA-seq expression of DEGs constituting solute carrier family genes in KO versus WT embryonic brains exposed to 30 ppm FA maternal diet. Scaled expression is shown as the Z-score of TPM values. (**D**), Number of up and down-regulated DEGs defined by dietary FA 30 ppm or 0.1 ppm vs 3 ppm pairwise comparisons of E11.5 embryonic brain samples per genotype. Differential cut- off was FDR-adjusted *P* value < 0.05. (**E**), Venn diagram of all down-regulated DEGs defined by KO vs WT exposed to 30 ppm FA in (A), 0.1 ppm vs 3 ppm in WT and HET groups in (B). The number of genes in the overlap is highlighted with green in bold. (**F**), GO analysis of the 71 overlapping down-regulated DEGs as defined in in (E). (**G**), Violin plot for the normalized expression level of the 71 overlapping genes in WT, HET and KO stratified by diets. (**H**), Integrative Genomics Viewer (IGV) snapshots of DMRs tracks identified by RRBS on *Slc10a4* of WT, HET, and KO stratified by diets (left). Locations of DMR from WGBS of KO vs WT antNPCs (*34*), CpG island (CpGi) annotation and TET1 ChIP-seq signals in EpiLC (*35*), are indicated in the bottom panel. Right, RNA-seq TPM expression of *Slc10a4* per group (right).

*Tet1*^-/-^ versus *Tet1*^+/+^ brains under control diet exhibited few DEGs, suggesting that loss of TET1 alone minimally influences gene expression in E11.5 brains, unlike in antNPCs (*34*). Puzzled by this observation, we examined RNA-seq IGV tracks over the *Tet1* gene locus and observed transcript expression in WT E11.5 brains, elevated from the low levels detected in antNPCs *in vitro* and consistent with the post-gastrulation re-activation of *Tet1* expression in the embryonic brain (*33*). However, expression of a somatic *Tet1* isoform from a downstream TSS in KO brains was also detectable at low levels (Supplementary Fig. 7B). Expression of this somatic short TET1 isoform, which is missing the N-terminus CXXC domain and exhibits reduced global chromatin binding compared to the full-length embryonic isoform, may be sufficient to rescue gene expression but not DNA hypermethylation at CpG island promoter regions where full-length TET1 predominantly occupies (*58*). The prevalence of hyperDMRs resulting from loss of *Tet1* under control diet, with minimal DEGs, suggest that elevated DNA methylation has largely silent effects on gene expression at E11.5, or may be persistent aberrations originating during an earlier stage in development.

Next, to identify DEGs resulting from abnormal maternal FA levels, we conducted pairwise comparisons between excess FA (30 ppm) and depleted FA (0.1 ppm) versus control (3 ppm), per genotype. Notably, FA deficiency exerted the most pronounced effect on gene expression in HET embryos, resulting in 265 up- and 777 down-regulated DEGs (FDR adjusted *p*-value < 0.05) (Fig. 6D). This highlights a synergistic impact between *Tet1* haploinsufficiency and folate depletion. FA deficiency also impacted WT brains with 66 up- and 373 down-regulated DEGs but had negligible effects in KO brains (Fig. 6D). In contrast, excess FA has a minor impact, resulting in only 28 down- regulated DEGs in KO brains. The predominantly loss of gene expression caused by FA depletion in WT embryos was associated with reduced global histone acetylation, in alignment with a loss of accessible chromatin, but not with global changes in the facultative heterochromatin mark histone H3 lysine 27 tri-methylation (Supplementary Fig. 7C). We also noted that genes related to folate-cycle homeostasis, including *Dhfr*, *Fpgs*, and *Ggh*, were not differentially expressed between different groups (data not shown).

Given that DEGs were observed primarily in three pairwise comparisons – KO vs WT in 30 ppm FA diet, 0.1 ppm vs 3 ppm in WT and 0.1 ppm vs 3 ppm HET embryos, we overlapped the composite set of DEGs to identify a subset common to all. From the Venn overlap, we identified 71 common DEGs (Fig. 6E). GO analysis of these 71 genes showed a robust enrichment of neurotransmission-related terms, including “neurotransmitter transport”, “vesicle-mediated transport in synapse”, and “synapse organization” (Fig. 6F). Analysis of normalized expression levels confirmed that these 71 genes indeed exhibited reduced expression selectively in KO embryos exposed to 30 ppm FA, but in all genotypes under 0.1 ppm FA (Fig. 6G). The loss of expression of neurotransmitter and synaptic function related genes aligns with the observed loss of phospholipid metabolites in KO embryos under excess FA. Moreover, we observed more DEGs in the HET brains under FA depletion, which coincided with a significant increase in the occurrence of brain malformation, and reduction in methylation potential compared to HET embryos under control diet (Fig. 4D,F). These observations suggest an excess of FA in the absence of TET1 may paradoxically mimic FA deficiency by a down- regulation of multiple solute carriers involved in neurotransmitter transport, resembling a compensatory response to reduce influx of FA. Interestingly, while hyper DMRs resulting from *Tet1* KO were prevalent across all diet groups, for instance at the gene promoters of solute carriers *Slc10a4* and *Slc32a1*, these hyper DMRs were associated with a significant loss of gene expression only under excess FA conditions (Fig. 6H and Supplementary Fig. 7D). These results strongly indicate a synergy between excess FA status and DNA hypermethylation leading to gene silencing. Moreover, either an excess of FA coupled with *Tet1* loss or FA deprivation impacts expression of a common set of DEGs, implicating intricate interactions between *Tet1* and folate 1C cycling in the regulation of multiple solute carriers involved in neurotransmitter transport.

Folate transport across cell membranes involves three distinct mechanisms, via receptor mediated endocytosis of the folate receptor alpha and beta (FRα, gene *Folr1;* FRβ, gene *Folr2*), via the reduced folate carrier (RFC, gene *Slc19a1*) in systemic tissues, and via the proton-coupled folate transporter (PCFT, gene *Slc46a1*) located primarily in the kidney, gastrointestinal tract, and the choroid plexus in the ventricles of the brain (*59*). The *Slc46a1* gene has a CpG island promoter known to be regulated by DNA methylation (*60*). TPM values of *Slc46a1*, *Slc19a1, Folr1* and *Folr2* in our RNA-seq analysis of 129S6.Cg E11.5 brains did not show any significant changes in expression of these genes due to differences in *Tet1* genotype and FA diet (Supplementary Fig. 8A). In CD1 E11.5 embryos, we observed modest down-regulation of *Slc46a1* in the KO brains under both excess and normal FA diet (Supplementary Fig. 8B). Nonetheless, our previous whole-genome bisulfite sequencing (WGBS) revealed hypermethylation at the *Slc46a1* promoter in *Tet1*^-/-^ antNPCs (which mimic the E8.5 neuroepithelium) (*34*) (Fig. 7A). This region is occupied by TET1 in primed epiblast-like cells, implicating *Slc46a1* as a direct TET1 target gene in early development pre-gastrulation. Therefore, we examined a potential regulation of *Slc46a1* by TET1 in post-gastrulation neuroectoderm cells prior to neural tube closure, which can be mimicked by antNPCs converted from ESCs by *in vitro* differentiation. Notably, *Slc46a1* expression and PCFT protein were decreased in 129S6.B6-*Tet1*^-/-^ antNPCs compared to *Tet1*^+/+^ cells, but not in cells with the F1 background (Fig. 7B, C), while *Slc19a1* was not affected (Fig. 7B). These results suggest that PCFT expression in neural tissues in early development is affected by TET1 specifically in an NTD- susceptible strain.

**Fig. 7.**
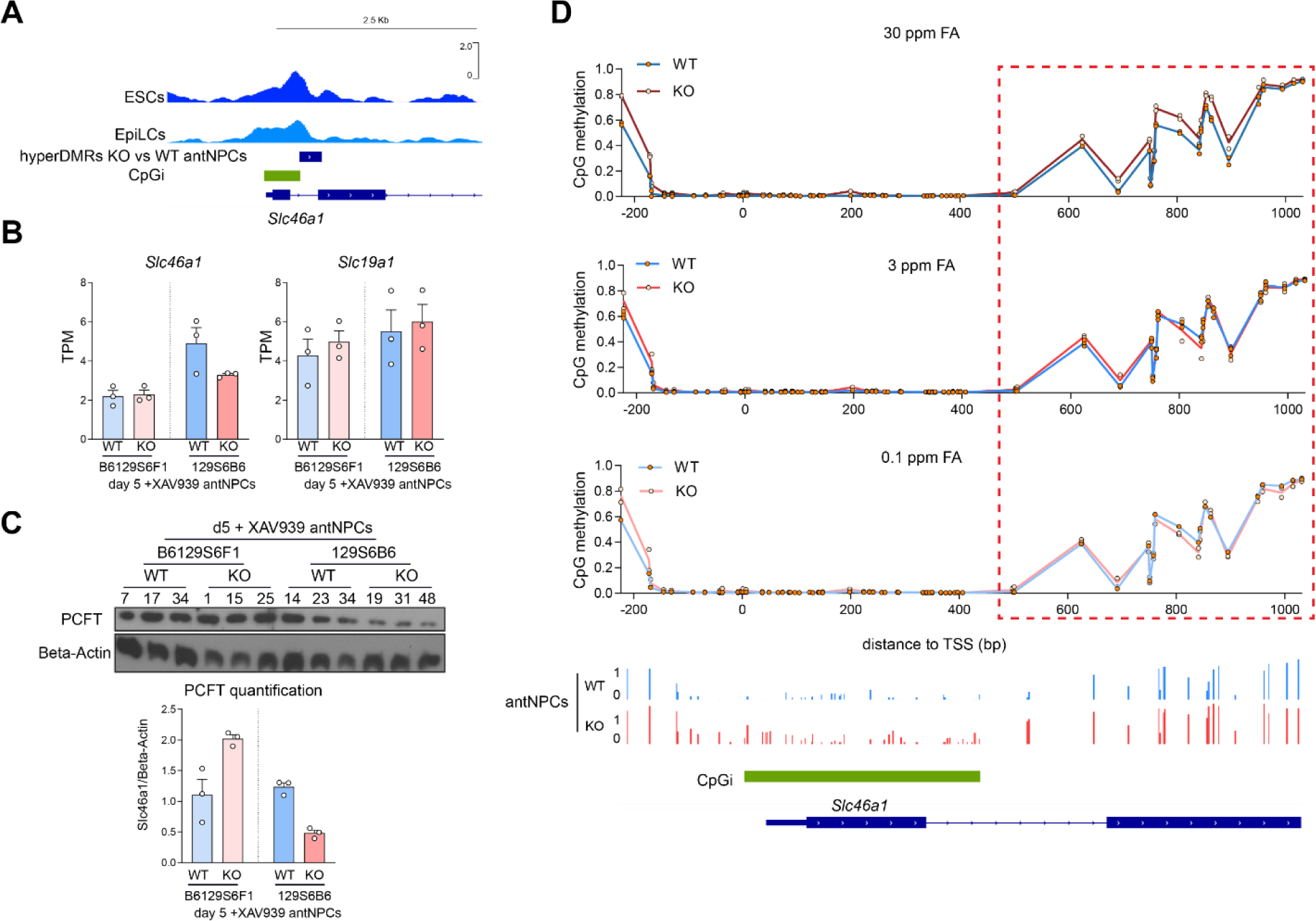
Excess FA coupled with *Tet1* loss results in hypermethylation on the promoter region of folate transporter *Slc46a1*. (**A**), IGV snapshots of TET1 ChIP-seq tracks in mESCs (*35*) and EpiLCs (*33*), illustrating the occupancy of *Tet1* on the promoter region of *Slc46a1*. HyperDMRs, resulting from loss of *Tet1* in antNPCs generated by 5 days of directed differentiation in the presence of a Wnt inhibitor (day 5 + XAV) (*34*). (**B**), RNA-seq expression of *Slc46a1* and *Slc19a1* in KO and WT antNPCs of F1 and 129S6.B6 strains. (**C**), Western blot of PCFT in KO and WT antNPCs of F1 and 129S6.B6 strains. ACTB, beta- actin loading control. Quantification plot is shown at the bottom panel. (**D**), Targeted amplicon bisulfite sequencing analysis of CpG methylation levels at the promoter region of *Slc46a1* in WT and KO whole embryos exposed to maternal diet of modified FA levels. X- axis 0 indicates position of the TSS. IGV tracks below show 5mC methylation levels in F1 WT and KO d5 + XAV antNPCs over the corresponding region (*34*).

Encouraged by these results, we asked whether dietary FA levels may modulate *Slc46a1* promoter methylation levels in *Tet1*^-/-^ embryos. Because the DMR at *Slc46a1* identified in KO versus WT antNPCs was not covered by RRBS, we conducted targeted bisulfite sequencing over the region to re-analyze whole embryos collected from dams on modified FA diets on the 129S6.Cg background (Supplementary Fig. 8C,D). Interestingly, FA excess drove an elevation in CpG methylation at these CpG shores in *Tet1*^-/-^ embryos; conversely, FA depletion caused slightly lower CpG methylation in KO compared to WT, whereas methylation differences were not discernible between WT and KO under control 3ppm FA diet (Fig. 7D). These results affirm that dietary folate status interacts with *Tet1* to regulate promoter DNA methylation of multiple membrane transporters, including a folate transporter expressed in the brain.

## Discussion

The complexities of gene-environmental influences in birth defect susceptibility pose a challenge to our comprehension of disease etiologies. In this study, we adopt an epigenetic perspective to elucidate how variations in both genetic makeup and environmental exposures may converge to influence the risk for developing NTDs. We demonstrate that the DNA dioxygenase TET1, a newly identified candidate gene for NTDs, is a crucial link connecting FA, 1C metabolism, DNA methylation and genetic modifiers influencing NTD risk. We identified a QTL for NTD susceptibility linked to the loss of TET1. The QTL peak harbors among prioritized candidates implicated in cellular trafficking and signaling pathways, aligning with a cell non-autonomous mechanism by which loss of *Tet1* function affects NTC, and strain-dependent sensitivities to developmental signaling. Further, we uncovered that excess FA status interacts with loss of *Tet1*, resulting in the down-regulation of membrane transporters and disruption of FA intake, providing a mechanistic basis for NTDs resistant to FA supplementation. Conversely, FA deficiency interacts with *Tet1* gene dosage, impacting cellular methylation potential and NTD susceptibility. Overall, the interplay between epigenetic regulation by TET1 and 1C metabolism driven by folate converges on phospholipid metabolism and expression of neurotransmitter genes, ultimately affecting neurodevelopment beyond NTC.

Strain-dependent variations in NTDs have been observed in various other mouse mutants, with 129 strains displaying higher disease susceptibility while B6 strains generally do not (*12, 13, 61*). To our knowledge, few studies have ventured into the exploration of modifier loci for these variations (*62–64*). Conventional gene knockout studies are frequently conducted in mice using congenic inbred strains, which may not adequately capture the complex genetic landscape associated with human NTDs. The stark disparity in NTD penetrance among different strains of *Tet1* mutants we generated, thus offers a unique opportunity to identify the genetic factors that modify NTD susceptibility. Our QTL analysis mapped the recessive risk locus to a 0.9 Mb region in the 129S6 genome on chromosome 9. Interestingly, this locus contained a missense variant in a gene, *Snx1*, previously known to cause NTDs when mutated together with *Snx2* (*65*). *Snx1* is part of the sorting nexin family and is required for proper endosome to Golgi-system trafficking via the retromer pathway (*66*), which is important for regulating signaling pathways, such as the EGF pathway (*67, 68*). Family members *Snx3* and *Snx13* have also been implicated in mouse NTDs (*45, 69*), suggesting that defects in endocytosis can contribute in the etiology of NTDs (*70*). Interestingly, *Snx3* plays a critical role in the intracellular trafficking of both canonical and non-canonical WNT proteins for secretion, and *Snx3*^-/-^ mice display 100% cranial NTDs (*45*). Possibly, the P117S variant forms a *de novo* phosphor-site, inhibiting SNX1 function. Dysfunction of SNX1 reduces the recycling of Wntless protein (*71, 72*), resulting in the reduced secretion of Wnt-ligands which are required for NTC (*47, 49, 71, 72*).

*Snx1*, the only NTD candidate gene within the 1Mb QTL peak, is also situated at a TAD boundary between two intron variants at *Csnk1g1* and *Dapk2*, raising the possibility that the strain variation may affect expression via changes in higher chromatin interactions. Interestingly, *Csnk1g1* is a member of the casein kinase I gene family, encoding a serine/threonine kinase known to couple the extracellular Wnt signal and the cytoplasmic signal transduction machinery by phosphorylating the Wnt receptor LDL-receptor-related protein 6 (LRP6) (*46*). SNX1 itself is a target of phosphorylation by DAPK2 (*73*). The observed strain-dependency in co-regulation of gene expression spanning the QTL suggests that modified expression of multiple interacting genes within the locus, rather than complete loss of expression of any particular one, may act together to cause the NTD phenotype. Our NTD-sensitive 129S6.B6 *Tet1* mutant strain still lacks complete penetrance; the 129S6 chr 9 QTL is associated with increased NTD risk, but by itself does not show an absolute causal link. These results suggest the presence of other NTD-modifying QTLs that require more genetically diverse mouse models for their identification.

In the Genome Aggregation (gnomAD) database, TET1 has a probability of being loss-of-function (LoF) intolerant (pLI) score of 1, indicating extreme intolerance for LoF in humans. In agreement, a recent study of 506 spina bifida subjects validated *TET1* as an ultra-rare deleterious variant (URDV)- containing NTD candidate gene(*74*). The *TET1* deleterious variant (p.E1372K) identified in this largely Caucasian cohort co-occurred with URDVs associated with WNT signaling and cell migration pathways. Thus, our strain-comparative analysis in *Tet1* null mouse model parallels observations in a human patient cohort, suggesting that *TET1* loss-of-function mutations, while causing cranial NTDs (exencephaly) in mice, may also contribute to caudal (spinal) NTDs in humans.

The most recent recommendation by the US Preventive Services Task Force re-affirmed the substantial NTD-protective benefit of FA supplementation, when taken at 0.4 to 0.8 mg daily during the peri-conceptual period (*75*). However, whether excessive maternal FA intake, as high as 5 mg/day in certain medical conditions such as those linked to low folate levels, may introduce aberrant DNA methylation events or *de novo* point mutations that pre-dispose the offspring to chronic diseases in adulthood is a concern that warrants further investigation (*23, 76, 77*). The influence of excessive maternal dietary folate on DNA methylation has been reported to affect specific loci including imprinted genes (*78–81*). In this study, we used 30 ppm FA as a dose representing a 10-fold surplus of FA in rodent diet, in accordance with other studies (*23, 54, 56*). Excess FA supplementation can cause teratogenic effects on fetal brain development and confer susceptibility to embryonic defects (*82–84*). While we observed mild teratogenic effects of excess FA inducing brain malformation (8.5%) during embryonic neurodevelopment, our genome-wide profiling also uncovered hundreds of DMRs predominantly with gains of DNA methylation in WT mouse embryos. The impact of excess maternal FA on the WT DNA methylome is generally not associated with gene expression changes by E11.5. It is also more subdued compared to FA deprivation, which induced DMRs more extensively with both gain and loss in DNA methylation affecting pathways associated with development and Wnt signaling, concomitantly with gene expression changes predominantly downwards. However, our study revealed adverse effects of excess FA accentuated in the absence of TET1.

While loss of TET1 would be expected to exacerbate DNA hypermethylation driven by excessive folate, we observed on the contrary a reduction in hyperDMRs, highly resembling the pattern seen in *Tet1* deficient embryos deprived of FA. This could be partly due to the limitation of RRBS coverage, which is biased towards CpG-rich regions. Previous WGBS studies by us and others have shown TET1 activities to localize to CpG island flanking regions (shores), where DNA demethylation serves to counter DNMT activities to preserve CpG island promoters in a fully unmethylated state (*34, 85*). Using targeted bisulfite sequencing at a CpG shore region at *Slc46a1*, not covered by RRBS, we showed that excess FA can indeed elevate DNA methylation levels in KO relative to WT embryos. Possibly, excess folate in the absence of TET1 activity may result in a re-distribution of DNA hypermethylation from CpG island to shore regions.

Alternatively, the reduction in TET1-dependent hyper DMRs by both an excess or depletion of FA may reflect a state of pseudo-FA deficiency in *Tet1* KO embryos exposed to excess FA. This is supported by the increased uptake of FA in WT and HET embryos exposed to excess FA, but the inability to do so in KO embryos. Transcriptionally, *Tet1* KO embryos under excess FA also failed to upregulate multiple membrane transporters compared to *Tet1* WT and HET. Further DEG analysis revealed an overlap with genes down-regulated by FA deficiency in WT and HET embryos, with a striking convergence on pathways associated with neurotransmitter transport and function.

Adverse effects of excess FA excess coupled with the loss of TET1 also resulted in reduced phospholipid metabolites and down-regulation of multiple membrane transporters associated with promoter DNA hypermethylation. Of note, PCs are the most abundant phospholipids in mammalian cellular membranes, and in general their synthesis requires methylation reactions involving phosphatidylethanolamine methyltransferase (PEMT) and betaine-homocysteine methyltransferase (BHMT), both dependent on folate cycling of one-carbon via S-adenosylmethionine (SAM) (*86–88*). The absence of *Tet1* may shift SAM utilization under excess folate cycling towards uncurbed DNMT activities at the expense of PC synthesis, ultimately affecting transport of neurotransmitters and synapse formation. In this regard, the loss of *Tet1* mimics the reduction of MTHFR protein expression and activity by excess FA, which creates a pseudo-MTHFR deficiency that results in lower SAM/SAH ratio and alterations on lipid metabolism (Christensen et al, 2015). These results underscore the need to evaluate high-dose maternal FA supplementation in the context of potential nutritional stressors that may disrupt cellular dioxygenase functions, particularly for adverse impact on fetal neurogenesis and post-natal neurological functions.

Despite folate bioavailability being a major nutritional modifier of NTD risk, mutations directly affecting folate-related genes are relatively rare in cases of mouse NTDs (*4, 13*). Of these, only a few, such as *Mthfd1l* (*89*), *Gldc* (*90*) and *Slc25a32* (*91*), result in NTDs under folate deprivation, and all are resistant to FA supplementation. Interestingly, supplementation with formate, a THF- independent donor, circumvents the metabolic blocks caused by these mutations and rescue the FA-resistance (*90, 91*). While over 300 mouse mutant models for NTDs exist in the literature, only a fraction have been characterized for responsiveness to FA supplementation (*92, 93*). Our detailed study of *Tet1* mutants unravels an epigenetic mechanism as a basis for FA resistance across strains. Possibly, *Tet1*^-/-^ embryos are not able to absorb FA due to down regulation of *Slc46a1*/PCFT, resulting in their non-responsiveness to FA supplementation (in 129S6.Cg and CD1 backgrounds). On the other hand, FA deprivation exposes an effect of *Tet1* haploinsufficiency on methylation potential and increased occurrence of congenital structural malformations including NTD. These observations suggest that the risk for birth defects associated with FA status may be modified by other environmental factors that affect TET activity and function, such as vitamin C (*94*). By this rationale, FA supplementation in combination with vitamin C , would offer further protective benefits against birth defects. By reversing aberrant DNA methylation by informed nutritional intervention, FA-resistant cases of NTDs may potentially be converted into FA-responsive ones.

In the mouse, NTC begins at E8.5, resulting in closure of the anterior and midbrain-hindbrain neuropores by E9.5 and the posterior neuropore by E10.5 (*3*). The experimental end-point in this study at E11.5 would not have captured fully the dynamics of metabolomic and transcriptomic changes due to diet and genotype interactions during NTC. Indeed, our transcriptome analysis of embryonic brain tissues hardly detected any DEGs caused by loss of TET1 in the control diet, but revealed persistent down-regulation of solute transporters as a result of an interaction between excess FA and loss of TET1. Yet, the genome-wide DNA methylome analysis adequately captured widespread DMRs that more likely accumulated from diet x genotype manipulations during NTC, in line with a long-lasting presence of DNA methylation changes. For practical reasons, this study of E11.5 stage embryos enabled us to access a broader range of structural birth defects in addition to NTCs and also accurately dissect brain tissues (forebrain, midbrain, and hindbrain) to obtain tissue-specific epigenome and transcriptome profiles. Insights from this study will be the basis for further investigations of *Tet1* genotype x FA diet interactions earlier during the initiation of NTC, and later during postnatal development.

In summary, our study unravels the intricate layers of gene-environmental interactions influencing NTD risk. By decoding the interplay of TET1 with the genome, metabolome, and epigenome, we shed light on genomic loci and mechanisms associated with FA responsiveness and NTD risk, implicating cellular trafficking and signaling pathways. Targeted intervention strategies tailored to rectify these pathways with individualized micronutrient supplementation plans may have a major impact in preventing FA-resistant NTDs, a major focus of public health concern in the era post-FA fortification.

## Materials and Methods

### Animal maintenance, mouse strain breeding and timed pregnancies

*Tet1*^tm1Koh^ mice were maintained on a 12-hour day/night cycle with grain-based ‘chow’ maintenance complete feed (Ssniff, Germany). Breeding pairs are fed Ssniff breeding complete feed. Both regular maintenance and breeding diet contain 7 ppm FA. Unless stated otherwise, mice in experimental timed mating were fed maintenance complete feed. Breeding is maintained by crossing heterozygous with wild-type mice to obtain heterozygous offspring for timed pregnancy experiments. Pups were genotyped after weaning by performing PCR using primers as listed in Table S1. All experimental procedures on mice have been reviewed and approved under project P101/2016 “The epigenome:genome interactions involving TET1 in neural tube closure” by the KU Leuven Ethical Committee for Animal Experimentation in compliance with the European Directive 2010/63/EU.

The *Tet1*^tm1Koh^ line was originally created as a C57Bl/6 (B6) congenic strain and henceforth maintained by regular backcrossing to C57Bl/6J obtained from Jackson Lab. To transfer the *Tet1*^tm1Koh^ allele to alternative genetic backgrounds, we outcrossed it (> 3 generations) to the outbred CD1 stock. Subsequently, we backcrossed B6-*Tet1*^tm1Koh^ mice to an inbred 129S6 (also known as 129SvEvTac) strain. We considered mice obtained from 5-6 generations of backcrossing as incipient congenic, and those after >10 generations as congenic. Wherever indicated in the text, the number of backcross generations is prefixed by the letter N.

To obtain embryos from timed pregnancies, *Tet1*^+/-^ mice were paired and monitored for 5 days, checking every morning for the presence of a copulation plug. The morning of detecting a copulation plug was considered to be E0.5. Pregnant dams were sacrificed at indicated time-points using cervical dislocation. Embryos were dissected from individual decidua in ice-cold PBS. Following excision of the yolk sac, embryos were imaged using Leica Application Suite V4, and assigned a preliminary score according to the EMAP definition by a researcher. These scores were subsequently validated by two or three independent researchers based on images. Exencephaly was scored as “NTD”. For FA-modulation experiments, we also observed other malformations, that were classified as hindbrain deformity, eye defects, gross malformation of multiple visceral organs, and growth delay based on somite counts. Embryo heads or headfolds were dissected if needed and snap frozen in liquid nitrogen or fixed in 4% PFA over night at 4°C. For embryos ≥ E9.5, a portion of the yolk sac, free of maternal blood, was collected for genotyping. Genotyping primers are listed in Table S1.

### FA modulation via the diet

To manipulate dietary FA levels in the mice, we obtained custom diet with modified FA concentrations. We designed three different FA diets: FA-excess (30 ppm), FA-control (3 ppm) and FA-depleted (0.1 ppm) in accordance with previous studies (*23, 54, 95*). To establish a well-defined level of maternal folate availability, custom diets were supplemented with 1% succinyl sulfathiazole (SST), an antibiotic that inhibits folate synthesis by intestinal flora, as used in previous studies (*54, 95*). In every experimental cohort, we adapted females for 4 weeks to the 3 groups of custom diets; littermate females were randomly assigned to different diet groups. Stud males were re-used in timed pregnancies on a rotational basis across all diets. To determine whether the addition of SST in our custom diets by itself affected NTD phenotypic penetrance, a group of CD1-*Tet1*^tm1Koh^ mice were adapted with customized diets containing 2.7 or 7 ppm FA (matching level in regular chow) supplemented with or without 1% SST. At experimental end-points, dams were euthanized for embryo collection. The sex and genotype of individual E11.5 embryos were determined by PCR analysis with isolated yolk sac tissues. For the collection of biological replicates in next-generation sequencing analyses, we selected a *Tet1*^-/-^ embryo matched by somite numbers to a littermate WT and/or *Tet1*^+/-^ embryo of the same sex in every replicate. For each downstream assay, we collected 4 to 6 replicates from independent litters, representative of the phenotypes observed and matching numbers of male and female replicates.

### ESC culture maintenance

ESCs were cultured as previously described (*34, 35*). All murine ESC lines were cultured on mitotically inactivated C57BL/6J mouse embryonic fibroblasts (MEFs) in standard ESC culture medium composed of knock out DMEM (Invitrogen, 10829-018), 15% ESC-qualified fetal bovine serum (Invitrogen), 2 mM L-glutamine (Invitrogen, 25030-024), 0.1 mM 2-mercaptoethanol (Invitrogen, 31350-010), and 100:100 units:µg/ml penicillin: streptomycin (Sigma, P4333; or Gibco, 15140122), supplemented with leukemia inhibitory factor (LIF) produced in-house as culture supernatant from a LIF over-expressing CHO-cell line. As feeder preparation, MEFs were cultured on 0.1% gelatin in MEF medium consisting of DMEM GlutaMAX (Invitrogen, 61965-026), 10% FBS (Sigma-Aldrich F7524), 2 mM L-glutamine, 1 mM sodium pyruvate, 0.1 mM each of nonessential amino acids, 0.1 mM 2-mercaptoethanol and 100 U: 100 μg/mL penicillin: streptomycin. All lines were passaged using 0.25% Trypsin-EDTA (Invitrogen, 25200072). For this study we used 3 previously established littermate pairs of male *Tet1*^-/-^ and *Tet1*^+/+^ B6129S6F1-*Tet1*^tm1Koh^ ESC lines derived from mouse blastocysts. Three additional pairs of *Tet1*^-/-^ and *Tet1*^+/+^ 129S6.B6-*Tet1*^tm1Koh^ male lines were derived in this study. All lines were genotyped for the risk-allele on chromosome 9 by Sanger sequencing of the gel-purified PCR amplicon product. Primers are listed in Table S1. 129S6.B6-*Tet1*^tm1Koh^ ESC lines were derived and cultured as previously described (*33, 35, 50*). Both gender and *Tet1*^tm1Koh^ were genotyped using PCR and 3 pairs of male *Tet1*^-/-^ and *Tet1*^+/+^ lines were selected and used for subsequent experiments.

### GFP labeling ESCs with lentivirus

GFP transducing lentivirus was prepared as described previously (*34, 96*). In brief, 2 x 10^6^ HEK293T cells were seeded in a 10 cm dish and transfected the following day with 9 µg pSin-GFP, 8 µg pCMV- dR8.2, and 1 μg of pCMV-VSV-G plasmids using FugeneHD (Promega, E2311) in Opti-MEM (Invitrogen, 31985070). One day after transfection, the culture medium was replaced with ESC medium supplemented with 0.01 ng/ml recombinant murine LIF. Lentivirus-containing medium was collected from HEK293T cells 48h, 72h, and 80h after transfection, filtered with a 0.45 µm filter and concentrated ∼45x using Vivaspen20 columns (Sartorius, VS2032).

For labeling ESCs with GFP, serum cultured ECSs were feeder-depleted by twice 30-minute plating on 0.1% gelatin-coated plates, and 50,000 cells per well were seeded on 0.1% gelatin-coated 12- well plates. The next day 50 µl of concentrated virus was added together with 5 µg/ml polybrene (Millipore, TR-1003-G) to the culture media, followed by fresh media replacement 24h later. Following two passages on feeders in regular ESC medium, the ∼80-90 percentile of GFP+ cells were FACS sorted using a Sony MA900, to select cells with similar multiplicity of infections. Following sorting, cells were expanded for 1 passage on feeders and regular ESC medium and frozen down for subsequent blastocyst injection experiments.

### Diploid and tetraploid blastocyst injection assays

ESCs were thawed and passaged two times on feeders in regular ESC medium supplemented with 1000 U/ml LIF (Peprotech). Cells were split 1:5 on the day before blastocyst injection and medium was refreshed 2h before collecting the cells. Tetraploid embryos were produced via electrofusion of 2-cell stage embryos isolated from CD1 mice (40V, 30µs with a 250 µm electrode chamber in 0.3M mannitol, CF-10B pulse generator, BLS Ltd). The embryos were removed from the electrode chamber and washed in M2 (Merck), passed through 3 drops of Advanced KSOM (Merck) and kept for 30 minutes in a drop of KSOM (Merck, 37°C, 5% CO2 incubator) after which the tetraploid embryos were selected and put in a new culture dish with KSOM for 2 days until blastocyst were developed. For diploid blastocyst injection experiments, normal CD1 blastocysts were used. 10-12 *Tet1*^-/-^ *GFP*^+^-129S6.B6 ESCs were microinjected into each blastocyst and embryos were transferred to the uterine horn of pseudo pregnant CD1 female mice. At embryonic day E12.5, the embryos were dissected and morphologically assessed. Blastocyst manipulation and implantation were performed by the Mouse Expertise Unit of KU Leuven MutaMouse Transgenic Core facility under project P178/2020 “Generatie van genetisch gemodificeerde muismodellen” approved by the KU Leuven Ethical Committee for Animal Experimentation.

### Anterior neural progenitor cell (antNPC) differentiation from serum ESCs

AntNPC differentiation was performed as previously described (*34, 35, 52, 53*). Feeder- depleted serum cultured ESCs were plated on gelatin-coated plates at 10,000 cells/cm^2^ (90,000 cells per well in a 6-well plate) in advanced N2B27 defined media composed of a 1:1 mix of 1:1 DMEM/F12 (Invitrogen, 12634-010) and Neurobasal medium (Invitrogen, 10888-022), supplemented with 0.5x B27 without vitamin A (Invitrogen, 12587-010), 0.5x N2 (Invitrogen, 17502-048), 2 mM L-glutamine (Invitrogen, 25030-024), 40 mg/ml BSA fraction V (Invitrogen, 15260037), 0.1 mM 2- mercaptoethanol (Invitrogen, 31350-010) and 100 U: 100 µg/ml penicillin: streptomycin (Sigma-Aldrich, P4333; or Gibco, 15140122). The medium was changed every day during differentiation, except on day 1. Media change on day 0 and 2 were supplemented with 10 ng/ml bFGF (Cat#100- 18C, Peprotech). On day 4, 1.5x amount of medium was added. The experimental end-point was day 5.

### Western blotting

Cells were washed with ice-cold PBS and lysed on plate in ice-cold RIPA buffer (50 mM Tris at pH 8.0, 150 mM NaCl, 0.2 mM EDTA, 1% NP-40, 0.5% sodium deoxycholate, 0.1% SDS, containing 1 mM phenylmethylsulfonyl fluoride, 0.5 mM DTT, phosphatase inhibitor cocktail 2 and 3 (Sigma-Aldrich, P5726 and P0044) and protease inhibitor cocktail (Roche, 11836153001)). Lysates were scraped off the plates and transferred to pre-cooled microcentrifuge tubes, incubated on ice for 30 minutes, subsequently passed through a 26-gauge (26-G) needle and clarified by centrifugation for 15 min at 16.000 r.c.f. at 4°C, after which supernatant was collected and stored at -80°C. Protein concentration was measured using Bradford assay in a 96-well microplate format. Whole cell lysate samples were prepared in 1x Laemmli sample buffer (62.5 mM Tris-HCl at pH 6.8, 2.5% SDS, 0.002% bromophenol blue, 5% β-mercaptoethanol, 10% glycerol) and boiled for 10 min at 95°C. 10 to 20 µg of protein was loaded on an 8% (for TET proteins) or 10% (other proteins) SDS– polyacrylamide gel, electrophoresed in 1X running buffer (25mM Tris, 192 mM glycine, 0.1% SDS) and then transferred to a PVDF membrane with transfer buffer (25 mM Tris, 192 mM glycine and 20% methanol; 0.1% SDS added when transferring TET proteins). Membranes were blocked with 5% non-fat dry-milk tris-buffered saline with 0.1% Tween-20 (TBS-T). For phospho-proteins, membranes were blocked in 5% BSA in TBS-T. Subsequently, primary antibodies were diluted in 5% milk or 5% BSA (depending on blocking) and incubated overnight at 4°C, followed by incubation with corresponding secondary antibodies conjugated to HRP diluted 1:5000 in TBS-T with 5% nonfat milk for 1 h at room temperature. The signal was detected using Clarity Western ECL substrate (Bio-Rad 1705060) on AGFA Curix 60 Film Processor. Primary antibodies used in this study are: anti-non-phospho(Ser33/37/Thr41) (active)-CTNNB1 (Cell Signaling Technology, 8814, 1:1,000), anti- (total) CTNNB1 (Biosciences, 610153, 1:1,000), anti-phospho(Ser465/467)-SMAD2 (Millipore, ab3849-I, 1:1,000) , anti-(total) SMAD2/3 (Cell Signaling Technology, 3102 1:1,000), anti-ACTB (Sigma-Aldrich, A1978, 1:2000).

### Mass spectrometry

E11.5 embryos were snap frozen as whole embryos and processed for high performance liquid chromatography coupled to tandem mass spectrometry (HPLC-MS/MS) analysis as previously described (*97*). For ratiometric metabolite profiling, polar metabolites were extracted in -70°C 80% methanol/water for untargeted metabolomic analysis using a platform comprised of an Agilent Model 1290 Infinity II liquid chromatography system coupled to an Agilent 6550 iFunnel time-of-flight MS analyzer. Comprehensive chromatography of metabolites utilized a combination of aqueous normal phase (ANP) chromatography on a 2.1 mm Diamond Hydride column, and reverse-phase C18 hydrophobic chromatography, using both positive- and negative-ion modes for MS-based ratiometric quantification of metabolites. Mobile phases for aqueous chromatography consisted of (A) 50% isopropanol, containing 0.025% acetic acid, and (B) 90% acetonitrile containing 5 mM ammonium acetate. To eliminate the interference of metal ions on chromatographic peak integrity and electrospray ionization, EDTA is added to the mobile phase at a final concentration of 5 μM and the following gradient applied for separation: 0–1.0 min, 99% B; 1.0–15.0 min, to 20% B; 15.0 to 29.0, 0% B; 29.1 to 37 min, 99% B. Raw data were analyzed using MassHunter Profinder 8.0 and MassProfiler Professional (MPP) 15.1 software (Agilent technologies). One-way and two-way analysis of variance (ANOVA) (P < 0.05) were performed to identify significant differences between groups. Our in-house compound structural identity database included over 938 metabolites in our in-house MassHunter PCDL manager 8.0 database (Agilent) comprised of molecules that span 1C metabolism, vitamin co-factor, amino acid, nucleotides, carbohydrates, lipids and energy pathways. A molecular formula generator (MFG) algorithm in MPP is used to generate and score empirical molecular formulae, based on a weighted consideration of monoisotopic mass accuracy, isotope abundance ratios, and spacing between isotope peaks. A tentative compound ID is assigned when PCDL database and MFG scores concurred for a given candidate molecule. Metabolite structures are assigned based on monoisotopic neutral masses (<5 ppm mass accuracy), and chromatographic retention times, with confirmation by MS/MS fragmentation pattern matching to reference standards (*98, 99*).

The untargeted metabolic profiling approach precluded accurate measurement of bioactive forms of FA, such as 5-methyltetrahydrofolate (5-MTHF) and tetrahydrofolate (THF). As a result, metabolites of these bioactive forms of FA were excluded from the metabolomic data. Non-mammalian metabolites, metabolites with a retention time < 1.3 min and/or no confidence on the ID assignment were also excluded. Chiral structure specifications were removed, and lipid names were standardized according to the Lipidomics Standard Initiative. Statistical analysis was then performed using 2-way ANOVA to identify significant interactions between the diet and genotype factors (P < 0.05). Significant metabolite values were scaled using Z-score and visualized using the pheatmap R package (v. 1.0.12) (*100*).

### Targeted amplicon bisulfite sequencing library

We assayed the *Slc46a1* promoter methylation status by designing 8 amplicons of 150-290 bp each, spanning 1320 bp of the promoter region, for analysis by targeted bisulfite sequencing, performed as described previously (*34*). In brief, genomic DNA (gDNA) was extracted using the Purelink genomic DNA Mini Kit (Invitrogen, K182001) from E11.5 brains. DNA quality was assessed with Nanodrop, confirming the absence of RNA contamination with a 0.8% agarose gel stained with SYBR Safe. For bisulfite conversion, 1.5 µg of gDNA was eluted in 15 µl of elution buffer from the EpiTect Fast DNA Bisulfite kit (Qiagen, 59824). A 20 µl PCR reaction was composed of 0.5 µl of bisulfite-converted gDNA input, 300 nM each of forward and reverse primers (containing P7 and P5 tails, respectively, Table S1), PlatinumTM Taq DNA polymerase High Fidelity (Invitrogen, 11304- 011) and PCR kit buffers. PCR products were gel-extracted using the PureLink Quick Gel Extraction kit (Invitrogen, K210012). Amplicon concentrations were quantified with the QubitTM dsDNA HS Assay kit (Invitrogen, Q32854) and diluted to 15 nM.

Amplicons per sample were pooled to a maximum concentration of 5 ng/µl and assessed for DNA quality using a fragment analyzer (Agilent) and the Qubit™ dsDNA HS Assay kit (Invitrogen, Q32854). Amplicon pools were converted into sequencing libraries by performing a secondary PCR to incorporate indexes and sequencing adapters. The PCR reaction included 9 µl of DNA, 0.5 µl of custom p7 primer (125 nM), 0.5 µl of custom p5 primer (125 nM), and 1x 10 µl Phusion® High Fidelity PCR master Mix with HF buffer (Biolabs new England M0531S). The thermocycler program was 94 °C for 30 sec; 15 cycles of 94 °C for 10 sec, 51 °C for 30 sec, 72 °C for 30 sec; and a final extension at 72 °C for 1 min. Custom primers were provided with a unique dual index for sample labeling. After purification using AMPure XP beads following the manufacturer’s protocol, the library’s final quality was analyzed with a fragment analyzer and then pooled equimolarly. The concentration of the final pool was measured using qPCR (Kapa SYBR fast, Roche, KK4600), diluted to 4 nM and loaded onto a NovaSeq for PE150 sequencing, aiming for a minimum of 200,000 reads per amplicon and an average of 350,000 reads. For data analysis, reads were processed using Trim Galore! (v0.6.7) for quality, adapter removal, and exclusion of reads shorter than 20 bp. Subsequently, Bismark (v0.23.1) aligned the trimmed reads to GENCODE mm10 GRCm38.p6, with a maximal insert size of 500 bp, followed by methylation extraction. Only CpGs with a minimum coverage of 1000x were retained for analysis and were visualized over the genomic locus using a custom R script (v4.0.3).

### Exome-seq library preparation

Genomic DNA (gDNA) was extracted from whole embryos using the Gentra Puregene Tissue kit (Qiagen, 15063), according to the manufacturers’ specifications with one modification: to eliminate the high concentration of RNA in mouse embryos, 1 ul of 100 mg/ml RNase A (Qiagen, 19101) was added after Proteinase K digestion at 55°C for a further one-hour incubation at 37°C to degrade all RNA. The gDNA was quantified using the Qubit 2.0 DNA HS Assay (ThermoFisher, Massachusetts, USA) and its quality was assessed with the Tapestation genomic DNA Assay (Agilent Technologies, California, USA). Exome capture was performed using the IDT xGen Exome Research Panel v2.0 in conjunction with library preparation using the SureSelectXT kit (Agilent Technologies, California, USA) following the manufacturer’s recommendations and incorporating Illumina® 8-nt dual-indices. Library quality and quantity were assessed using the Qubit 2.0 DNA HS Assay (ThermoFisher, Massachusetts, USA), Tapestation High Sensitivity D1000 Assay (Agilent Technologies, CA, USA), and QuantStudio® 5 System (Applied Biosystems, USA). Equimolar pooling of libraries was performed based on QC values, and the libraries were then sequenced PE150 on an Illumina® Novaseq S4 flow cell (Illumina, California, USA) for 26 million reads per library, aiming for an approximate 100x coverage per library.

### Reduced representation bisulfite sequencing (RRBS) library preparation

Genomic DNA (gDNA) was extracted using the Purelink genomic DNA Mini Kit (Invitrogen, K182001) from E11.5 brains. DNA quality was assessed with Nanodrop, and absence of RNA contamination was confirmed with a 0.8% agarose gel stained with SYBR Safe. RRBS libraries were prepared to include (oxidative) bisulfite conversion using the Ovation RRBS Methyl-Seq kit (Tecan/Nugen, 0553-32), according to the manufacturer’s instructions. Per sample, 100 ng of gDNA as input was processed through oxBS workflow. To increase the coverage, we performed double digestion with both MspI (Supplied with the Nugen Kit), and TaqI (NEB, R0149S) with each sample. We first performed the MspI digestion at 37°C for 60 min, followed by the TaqI digestion at 65°C for another 60 min. After digestion, each library was ligated to custom single-indexed sequencing adapter (8 bp barcodes from the 96-Plex Adaptor Plate, Tecan, S02223) followed by final repair. Before oxidation and bisulfite conversion, libraries were purified and washed 3× with 80% acetonitrile to get rid of any residual ethanol, followed by incubation at 37°C for 5 min with denaturing buffer provided with the kit to denature the DNA. Libraries were oxidized by the addition of TrueMethyl oxidant solution incubating at 40°C for 10 min. Immediately after the oxidation, each library was converted with the bisulfite reagent solution provided with the kit, following the bisulfite conversion thermal cycler program in the instruction. Bisulfite converted DNA was then desulfonated with the provided desulfonation buffer at RT for 5 min, following 2× washing with 70% ethanol. For each library, the optimal amplification was optimized using qPCR; 1/5th of the libraries was added to amplification master mix containing SYBR Green and run on an Applied Biosystems StepOnePlus Real-Time PCR system for 30 cycles. Relative log-fluorescence vs amplification cycle was plotted out to manually determine the appropriate amplification cycles, selected within the middle to late exponential phase of amplification. Libraries were amplified between 8 and 13 cycles. Following amplification, libraries were purified with a final 1× clean-up using AMPure (Agencourt) beads. The quality of the libraries was assessed using an Agilent Bioanalyzer 2100 with the High Sensitivity DNA analysis kit (Agilent, 5067-4626) and concentration was determined using Qubit™ dsDNA HS Assay kit (Invitrogen, Q32854). The libraries were sequenced obtain minimally 20 million PE150 reads.

### RNA-seq library preparation

Total RNA was extracted from cultured cells using Trizol using the manufacturer’s instructions. RNA sequencing (RNA-seq) libraries were prepared from 4 µg of total RNA using the KAPA stranded mRNA-seq kit (Roche, KK8421) according to manufacturer’s specifications. 100 nM KAPA-single index adapters (Roche, KK8700d) were added to A-tailed cDNA, and libraries were amplified for 10 cycles. From embryonic brain tissues, RNA was extracted using RNeasy plus mini kit (Qiagen, 74136) according to the manufacturer’s instructions. The integrity of RNA was confirmed using Agilent Bioanalyzer 2100 with RNA 600 Nano kit (Agilent, 5067-1511). Libraries were prepared from 1 µg of total RNA using the KAPA Stranded mRNA Hyperprep Kit (96rxn) (Roche, KK8581) according to manufacturer’s specifications. 7 µM KAPA-single index adapters (Roche, KK8700d) were added to A-tailed cDNA, and libraries were amplified for 10 cycles. Finally, 1x library clean-up was performed using Agencourt AMPure XP beads (Beckman Coulter, A63881). Library fragment size was assessed using Agilent Bioanalyzer 2100 with the High Sensitivity DNA analysis kit (Agilent, 5067-4626) and concentration was determined using Qubit™ dsDNA HS Assay kit (Invitrogen, Q32854). Each library was diluted to 4 nM and pooled for sequencing on an Illumina Hiseq4000, aiming at 15-20 million SE50 reads per sample (19 million reads on average).

### scRNA-seq 10xGenomics library preparation

For single-cell library preparations, cells were washed with PBS and incubated for 5 minutes with TrypLE, which was subsequently quenched with advanced DMEM/F12. Single cells were washed 3x with 0.4% BSA in PBS. Cells were counted using the LUNA counter and an AO/PI fluorescent stain. Cells were loaded to target 5000 following the 10xGenomics protocol for single cell 3’ prime reagent kit v3.1 (single indexes). Quality control was performed and pooled according to instructions and sequenced following the 10xGenomics guidelines at 28-8-0-91 on a NovaSeq 6000 Instrument. First a shallow sequencing run was performed to assess quality of the libraries and estimate the number of captured cells, followed by a deeper sequencing run to reach a mean of ∼20,000 reads per cell. In total we captured 24,096 high quality cells after filtering, with on average 6,024 cells per sample.

### Exome sequencing analysis

FastQC was used for sequencing data quality analysis. Mapping, aligning, sorting, duplicate marking, and haplotype variant calling were performed using the Illumina DRAGEN Bio-IT Platform. The output resulted in vcf files. Subsequent filtering of genetic variants was conducted based on the following criteria: alternate allele count >= 0.2, max-missing 0.9 (allowing up to 10% missing data), minQ 30 (retaining sites with a Quality value above 30), min-meanDP 10 (ensuring a minimum mean depth of 10 for each site), and minDP 10 (setting the minimum depth for a genotype; individuals failing this threshold were marked as having a missing genotype). Following variant filtering, we utilized GATK to combine variants from all samples into a joint vcf file. We employed the vcf2gwas package (*101*) for genome-wide association analysis (GWAS).The Manhattan plot was generated using the R package qqman, with the top gene names marked on the peaks. The versions of software used were as follows: FastQC V0.11.8, SAMtools v1.9, GATK/Picard v4.1.6.0, VCFtools v0.1.17, Beagle V5.3, Vcf2gwas v0.8.8, and the mouse reference genome: mm10.

### Hi-C analysis

Publicly available Hi-C data from mouse embryonic stem cells (mESCs), neural progenitor cells (NPCs) and cortical neurons (CNs) (*39*) were downloaded in the .hic file format. The readHic function from the plotGardener (*102*) R package v. 1.4.1 was used to read the Hi-C contact data from .hic files. The triangular Hi-C interaction matrices were plotted using the plotHicTriangle function of the same package.

### RNA-sequencing analysis

Adapters, polyA/T tails, and bad quality reads (Phred score > 20) were trimmed using Trim Galore! (v0.6.4_dev) with default parameters. Reads were aligned to the transcriptome and quantified using Salmon (v0.14.1) (*103*) with default parameters using GENCODE release 23 of the mouse reference transcriptome sequences and the comprehensive gene annotation. Subsequently, the counts were imported into R (v4.0.2) using tximport (v1.18.0). Differentially expressed genes were defined using DEseq2 (v1.30.0) (*104*) (FDR adjusted p.val < 0.05) and log fold changes corrected using “ashr” method (*105*). GO term enrichment was performed using Cluster Profiler (3.18.1). Genes annotated to specific GO terms were extracted from ENSEMBLE version 98 using biomaRt (v2.56.1). TPM values were calculated using tximport.

### RRBS analysis

Processing of the raw reads was done according to the instructions of RRBS-library preparation kit (Tecan/Nugen). Reads were first trimmed for quality and presence of the adaptor using TrimGalore (v0.6.10) with the following command: “`trim_galore --paired -a AGATCGGAAGAGC -a2 AAATCAAAAAAAC`”, followed by removal of the diversity-adaptors using a custom Python script provided by Tecan/Nugen (“-b” flag is specified due to double digestion with both MspI and TaqI).

Subsequently, the trimmed reads were aligned to GENCODE mm10GRCm38.p6 using Bismark (v0.24.1) (*106*) with a maximum insert size of 500 bp. The Tecan/Nugen kit implements a UMI as the last 6 bp of I1 read, thus the aligned reads were deduplicated using UMI-tools (*107*), followed by methylation extraction with the bismark methylation-extractor (v0.24.1). On average, we reached a coverage of 8.7X and detected 2.6*106 CpGs with a minimal coverage of 5X per library, covering roughly 13% of mm10.

Differentially methylated regions (DMRs) were called based on two different strategies (Mulholland et al 2020) for every pair-wise comparison using MethylKit (v1.26.0) in R (v4.3.1), based on previous publication (*108*). First, differentially methylated CpGs were called (q-value < 0.05, Δ5mC < 10%) with a minimal coverage of 5X and were merged together (distance ≤ 250 bp, width ≥ 100 bp). Secondly, differential methylation scores (q-value < 0.05, Δ5mC < 10%) were calculated for covered regions, based on the UCSC annotation of promoters, CpG islands, CpGi shores, CpGi shelves and 1kb tiles with a 500 bp sliding window, containing at least 3 measured CpGs. From both strategies, regions that were overlapping were merged and filtered for containing minimally ≥ 2 differentially methylated CpGs and separated between hyper and hypo methylated DMRs. Per pairwise comparison, we only statistically tested CpGs that have an overall standard deviation of 2% 5mC within all samples (*108*). DMRs were annotated using AnnotatR (v1.26.0) and ChIPseeker (v1.36.0). Genes were associated with DMRs using the rGREAT (v2.2.0) package and GO term enrichment was performed using Cluster Profiler (4.8.1). DMR subsets were made using bedtools (v2.31.0). Profile plots were made using DeepTools (v3.5.5).

### scRNA-seq analysis

Reads were processed and aligned using Cell Ranger (v3.0.2) against mm10. The count matrix was imported in Seurat (v4.0.1) (*109*). Cells were selected based on criteria of 2000 to 8000 detected features per cell and a mitochondrial content between 1 and 10% as good quality cells. The cells are normalized per sample using the NormalizeData function in Seurat, using the “LogNormalize” method and a scaling factor of a 1000. The samples are integrated using the “fastMNN” method. The first 50 dimensions from the MNN reduction was used to determine the UMAP reduction and find clusters. These clusters were used in the “findAllMarkers” function to find genes that are differentially expressed between clusters.

## Supporting information

Data_S1_RNAseq_GO terms

## Acknowledgments

RNA-seq, 10xGenomics libraries were sequenced by Genomics Core Leuven (UZ Leuven). We thank Wouter Bossuyt, David Carbonez and Annelien Verfaillie at the Core for their excellent technical support in NGS experimental design and bioinformatic analysis. RRBS libraries were sequenced by Genome Scan (Leiden, the Netherlands). WES libraries were prepared and sequenced by Admera Health (NJ, USA). We thank Dan Gatti for guidance in the QTL analysis, and Paula Andrea Pimienta Ramirez for technical assistance in performing SNaPshot multiplex genotyping. We thank Sarah Cornet and Kelly Nguyen, our internship students, for their help in collecting the amplicons for *Slc46a1* bisulfite amplicon sequencing.

## Author contributions

K.P.K and B.K.V. designed and conceived the study. K.P.K. directed and supervised the study. K.P.K., B.K.V., and L.C. interpreted the data. L.Y. performed exome-seq and together with R.H.F. provided expertise on QTL analysis. R.H.F. and R.C. provided expertise and guidance on FA related experiments. B.K.V. and L.C. performed or supervised all experiments and molecular analysis techniques. S.C.T generated the Hi-C heatmaps based on the publicly available datasets, analyzed and interpreted the mass spectrometry data and generated the SLC gene expression heatmap. W.B. and M.S. maintained mice colonies. W.B. collected all samples for WES. Q.C. and S.S.G. performed mass spectrometry measurement of embryos and helped interpreting data. H.K. and R.K. collected embryos, performed histology, and defined the phenotype of Tet1tm1 embryos. B.K.V. and L.C. analyzed all data. K.P.K, B.K.V, and L.C. wrote the manuscript and prepared the figures.

## Data availability

All sequencing data will be deposited in GEO data repository. All other data are available in the main text or the supplementary materials.

