## supplementary figures and tables for "Epigenetic regulation by TET1 in gene-environmental interactions influencing susceptibility to congenital malformations"

**This file includes:**

Figs. S1 to S8

Tables S1 to S2

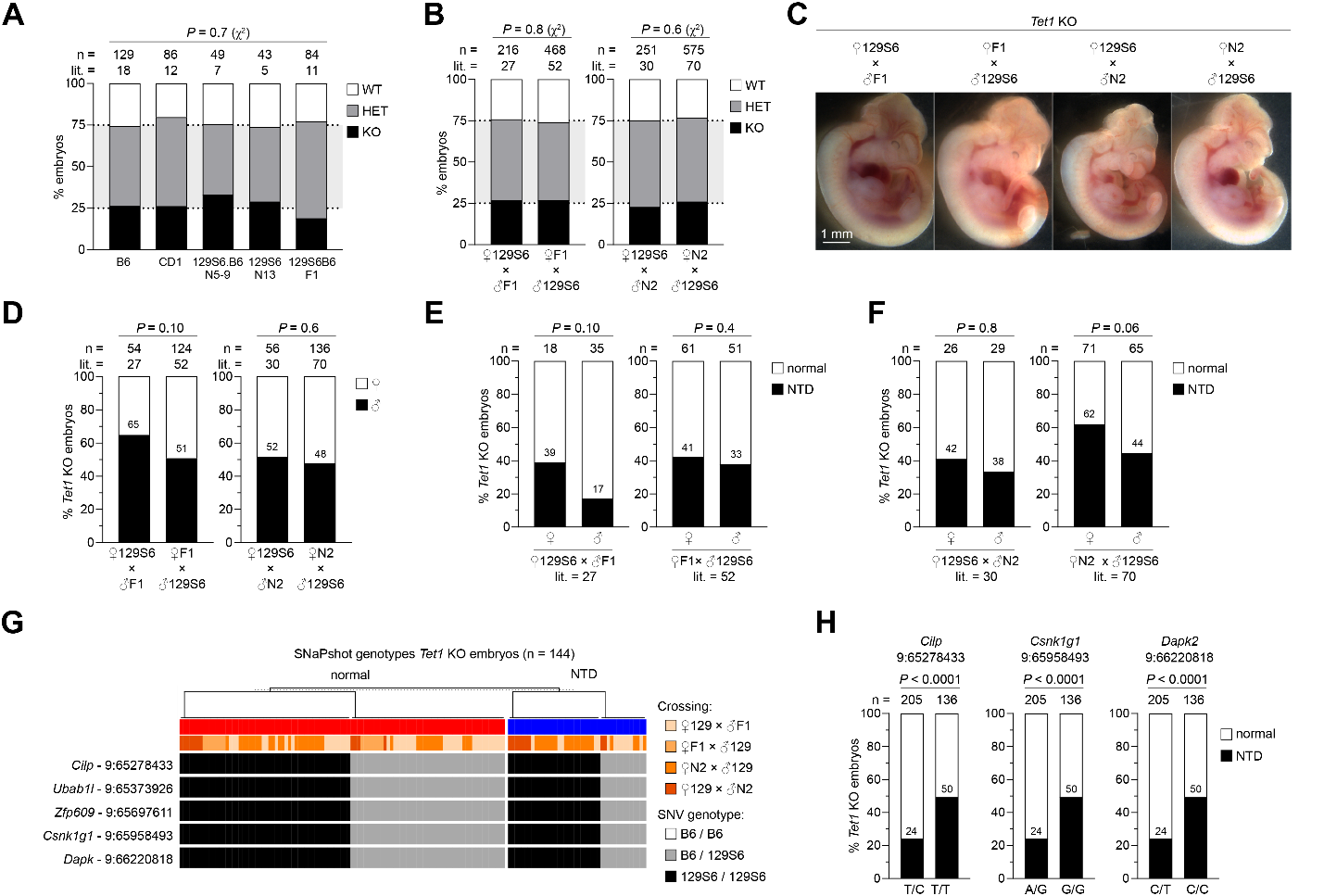

Fig. S1. Genotyping and sex ratios of NTD-resistant and NTD-susceptible strains of *Tet1*^-/-^ mouse embryos and SNaPshot multiplex PCR analysis at a chromosome 9 risk locus (related to main Fig.1).

(**A**), Proportion of *Tet1* genotype (WT=*Tet1*^+/+^, HET=*Tet1*^+/-^, KO=*Tet1*^-/-^) among offspring of heterozygous intercrosses on different genetic backgrounds. (**B**), Proportion of genotype in offspring from 129S6 x F1 or N2 (maternal x paternal) crossings and vice versa. Overall significance is determined using Chi square test in (A) and (B). (**C**), Images of *Tet1*^-/-^ embryos exhibiting NTDs from 129S6 x F1 or N2 crossings and vice versa. (**D**), Proportion of sex in all *Tet1*^-/-^ embryos collected from 129S6 x F1 or N2 crossings and vice versa. Number of male embryos are indicated above the black bars. (**E**) and (**F**), NTD rates in *Tet1*^-/-^ embryos obtained from 129S6 x F1 crosses and vice versa (E), and from 129S6 x N2 crosses and vice versa (F) stratified by sex. (**G**), Heatmap representation of SNaPshot multiplex PCR genotyping of 5 SNVs at the *Cilp*-*Dapk2* locus in normal and NTD-affected *Tet1* KO embryos from all four groups of strain intercrossing. (**H**), Allele effect plots of *Cilp, Csnk1g1 and Dapk2* within the most significant QTL peak region in the combined cohort of *Tet1* KO embryos sequenced by exome-seq and SNaPshot multiplex PCR. Numbers of NTD-affected embryos are indicated above the black bars in (E), (F) and (H). *P* values are calculated by Fisher’s Exact test in (D), (E), (F) and (H).

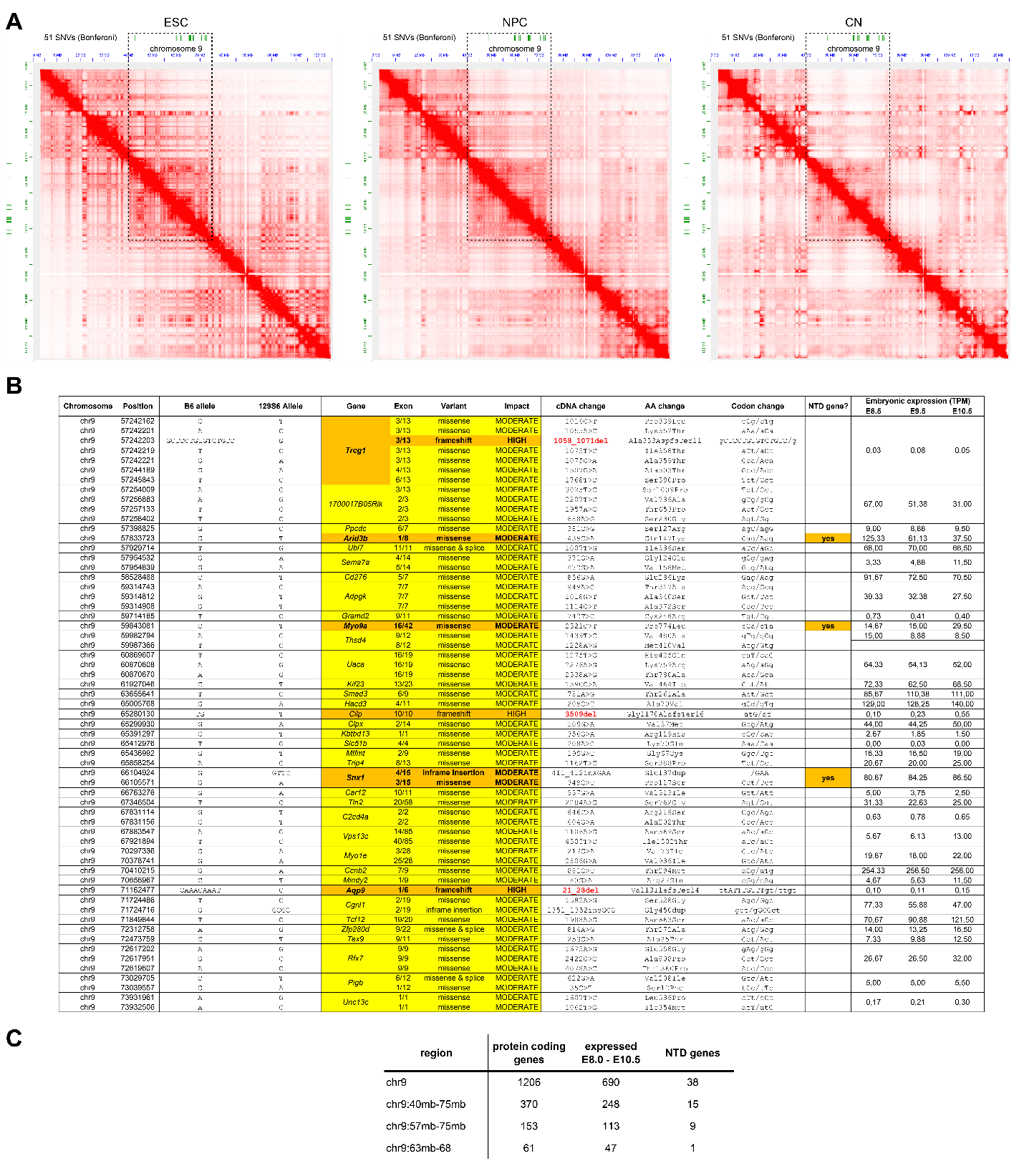

Fig. S2. Candidate modifier variants, associated genes and higher chromatin interactions at the QTL (related to main Fig. 2).

(**A**), Hi-C read densities in 10kb bins over chromosome 9 in ESCs (left), NPCs (center), or cortical neurons (CN, right). (**B**), Table of the 61 variants with a “high” or “moderate” predicted impact on protein function within the chr9:63mb-75mb locus. The chr9:63mb-68 QTL is indicated. (**C**), Number of protein coding genes on chromosome 9 filtered for expression during E8.5-E10.5 and implicated in NTD in the literature.

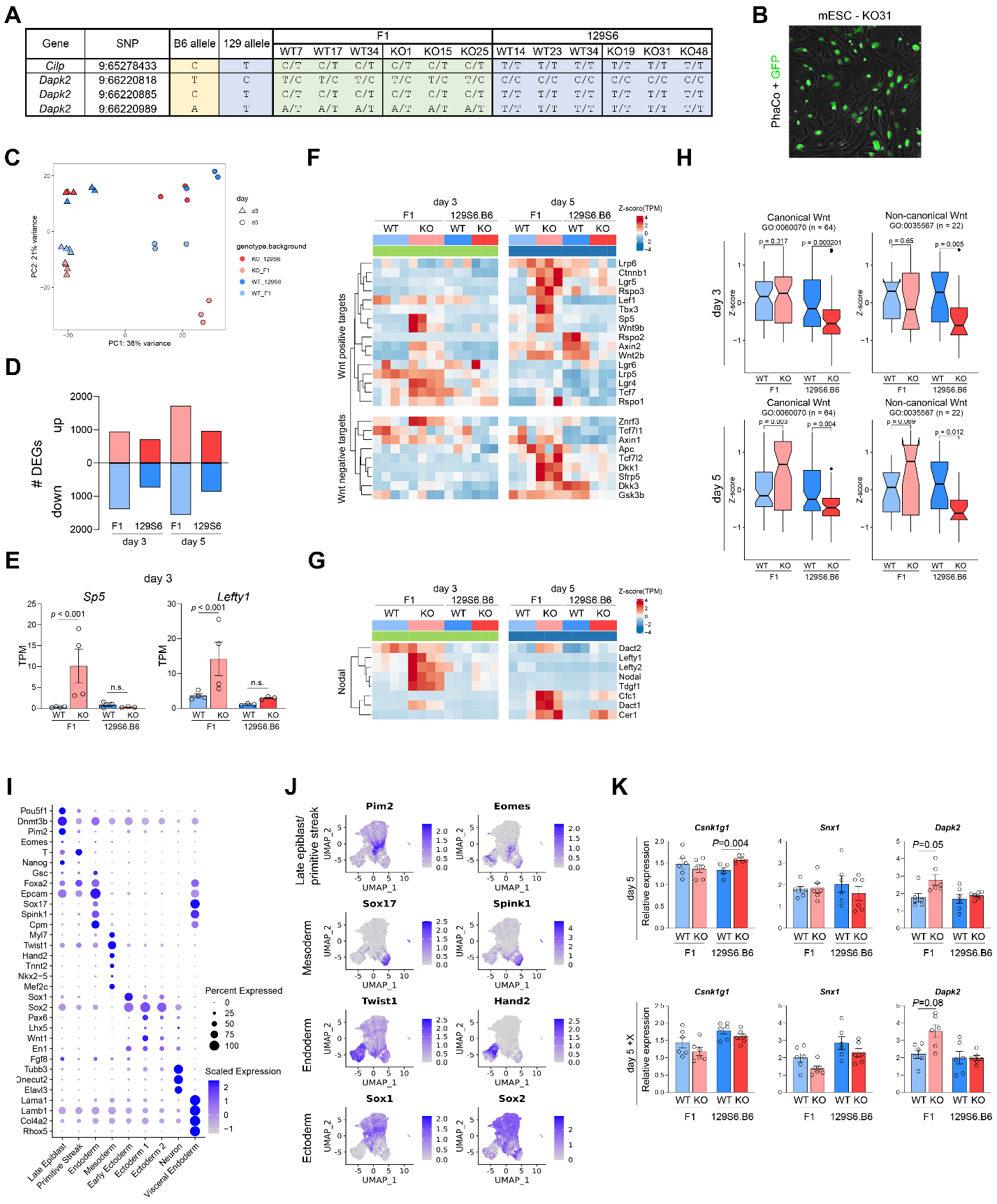

Fig. S3. Strain differences in sensitivity to developmental signaling recapitulated by NTD-resistant and -sensitive strains of ESC lines (related to main Fig. 3).

(**A**), Genotype of *Cilp* and *Dapk2* SNVs verified by clonal Sanger sequencing in each cell line generated from F1 and 129S6.B6 strains, *n*=3 per strain of *Tet1*^+/+^ and *Tet1*^-/-^. (**B**), Fluorescent image of the GFP^+^ 129S6.B6- *Tet1*^-/-^ ESC line used for blastocyst injection. (**C**), PCA plot for mRNA-seq of F1 and 129S6.B6 *Tet1*^+/+^ and *Tet1*^-/-^ cells on day 3 and day 5 of non-directed *in vitro* differentiation in neurobasal media. (**D**), Number of differentially expressed genes (FDR adjusted p-value < 0.05) per strain and differentiation time-point. (**E**), Gene expression of *Sp5* and *Lefty1* in RNA-seq transcripts per million (TPM) on day 3 of differentiation. (**F**) and (**G**), Heatmap of RNA-seq expression Z-score of genes involved in canonical Wnt (E) or Nodal signaling (F), on day 3 and day 5 of differentiation. (**H**), Boxplot showing the RNA-seq expression Z-scores of genes constituting the GO-term in Canonical Wnt signaling (left) and non-canonical Wnt signaling (right) in two strains of *Tet1*^+/+^ and *Tet1*^-/-^ cells, F1 and 129S6.B6 backgrounds, on day 3 and day 5 of differentiation. Number of genes per GO term is indicated in parentheses. The box represents the interquartile range (IQR), with the bottom and top edges corresponding to the first quartile (Q1, 25%) and the third quartile (Q3, 75%), respectively. The line inside the box represents the median (Q2, 50%) of the data. Whiskers extend from the box to a maximum of 1.5 times the IQR, and individual data points beyond the whiskers are considered outliers. (**I**), Dot plot of scRNA-seq expression of indicated marker genes to verify lineage identity of UMAP clusters determined on day 5 of differentiation, as shown in main figure 3(I)-(K.) (**J**), Heatmap of scaled expression of lineage markers for the indicated clusters on day 5 of differentiation. (**K**), Quantitative PCR measurement of for *Csnk1g1*, *Snx1*, and *Dapk2* gene expression on day 5 of differentiation. d5+X = d5 plus XAV-939, a Wnt inhibitor added on day 2. Relative expression is normalized to GAPDH. n=6 per genotype per cell line. *P* values are calculated by Student’s t-test.

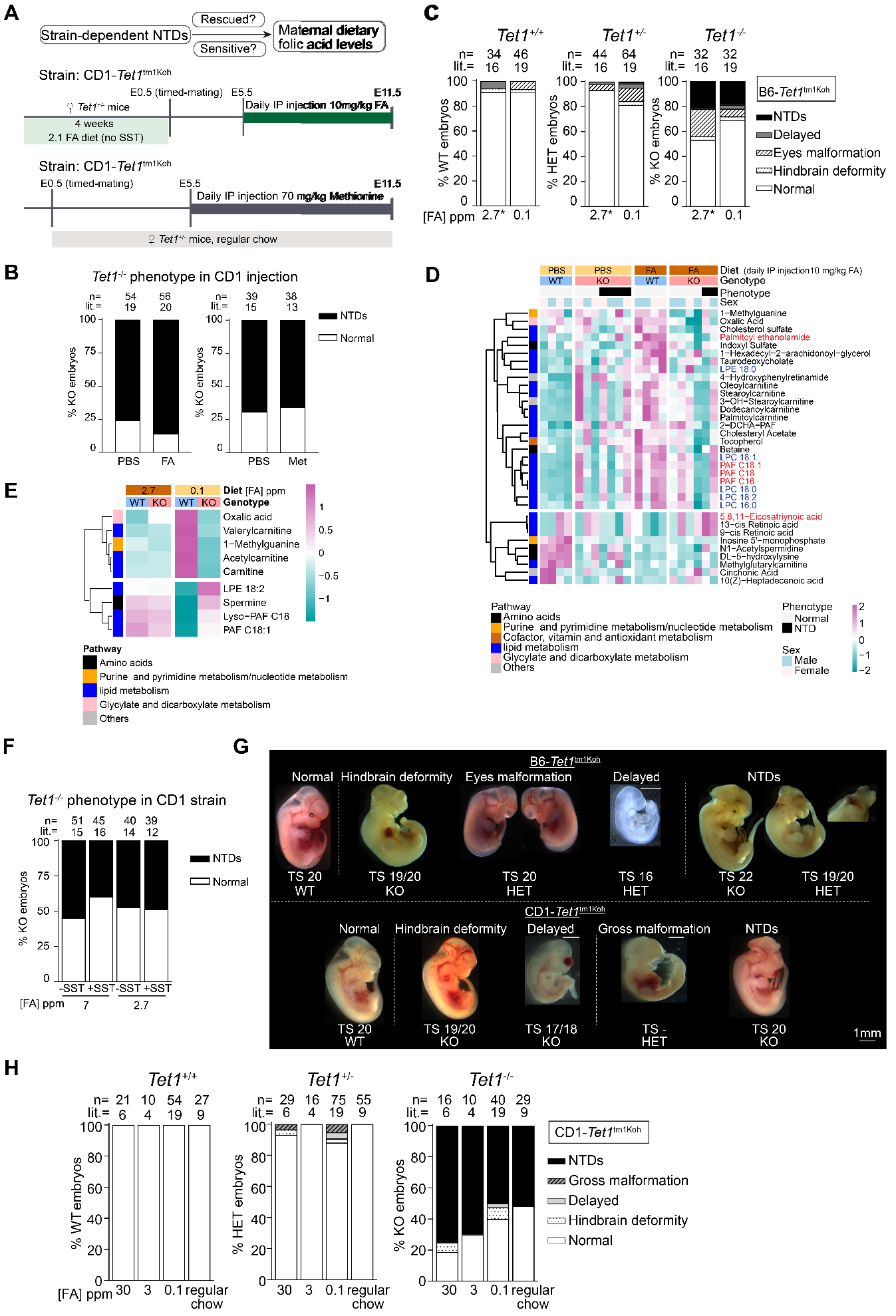

Fig. S4. Phenotypic and metabolomic analysis of CD1 and B6 embryos in response to maternal dietary FA excess or depletion, interacting with the loss of *Tet1* (related to main Fig. 4).

(**A**), Schematic (top) representation of the experimental designs to conduct FA rescue in CD1-*Tet1*^tm1Koh^ and FA depletion in B6-*Tet1*^tm1Koh^ mice. Protocols for daily intraperitoneal (IP) injection of 10 mg/kg FA in CD1-*Tet1*^tm1Koh^ heterozygous dams from E5.5 to E11.5 after adaptation to custom diet containing 2.7 ppm FA (middle), and 70 mg/kg methionine injection in dams maintained under standard chow containing 7 ppm FA (bottom). PBS was injected as vehicle control. (**B**), NTD rates in *Tet1*^-/-^ embryos of the CD1-*Tet1*^tm1Koh^ strain, in response to IP injection of FA and methionine. (**C**), Rates of phenotypes in *Tet1*^+/+^, *Tet1*^+/-^ and *Tet1*^-/-^ embryos of the B6-*Tet1*^tm1Koh^ strain exposed to FA-depleted maternal diet. (**D**), Heatmap of metabolites with significant changes (two-way ANOVA *p* < 0.05) in HPLC-MS/MS ion abundance detected in E11.5 whole embryos of CD1-*Tet1*^tm1Koh^ strain, in response to surplus FA injection. n=4, WT embryos in PBS and FA-injected groups; n=7, KO embryos in PBS-injected; and n=6, KO embryos in FA-injected groups. (**E**), Heatmap of representative metabolites in E11.5 whole embryo of B6-*Tet1*^tm1Koh^ strain under FA depletion, highlighting those with significant changes also in CD1-*Tet1*^tm1Koh^ embryos with surplus FA injection. n=4, WT embryos in 2.7 ppm group; n=5, WT embryos in 0.1ppm group; n=8, KO embryos in 2.7 ppm group; and n=5, KO embryos in 0.1ppm group. Two-way ANOVA *p* < 0.05. (**F**), Rates of NTDs in *Tet1*^-/-^ embryos of the CD1-*Tet1*^tm1Koh^ strain, in response to modified maternal diet containing either 7 or 2.7 ppm FA with or without SST. (**G**), Representative images of embryos with distinct phenotypes in B6-*Tet1*^tm1Koh^ and CD1-*Tet1*^tm1Koh^ strains. (**H**), Rates of phenotypes in *Tet1*^+/+^ *Tet1*^+/-^ and *Tet1*^-/-^ embryos of the CD1-*Tet1*^tm1Koh^ strain.

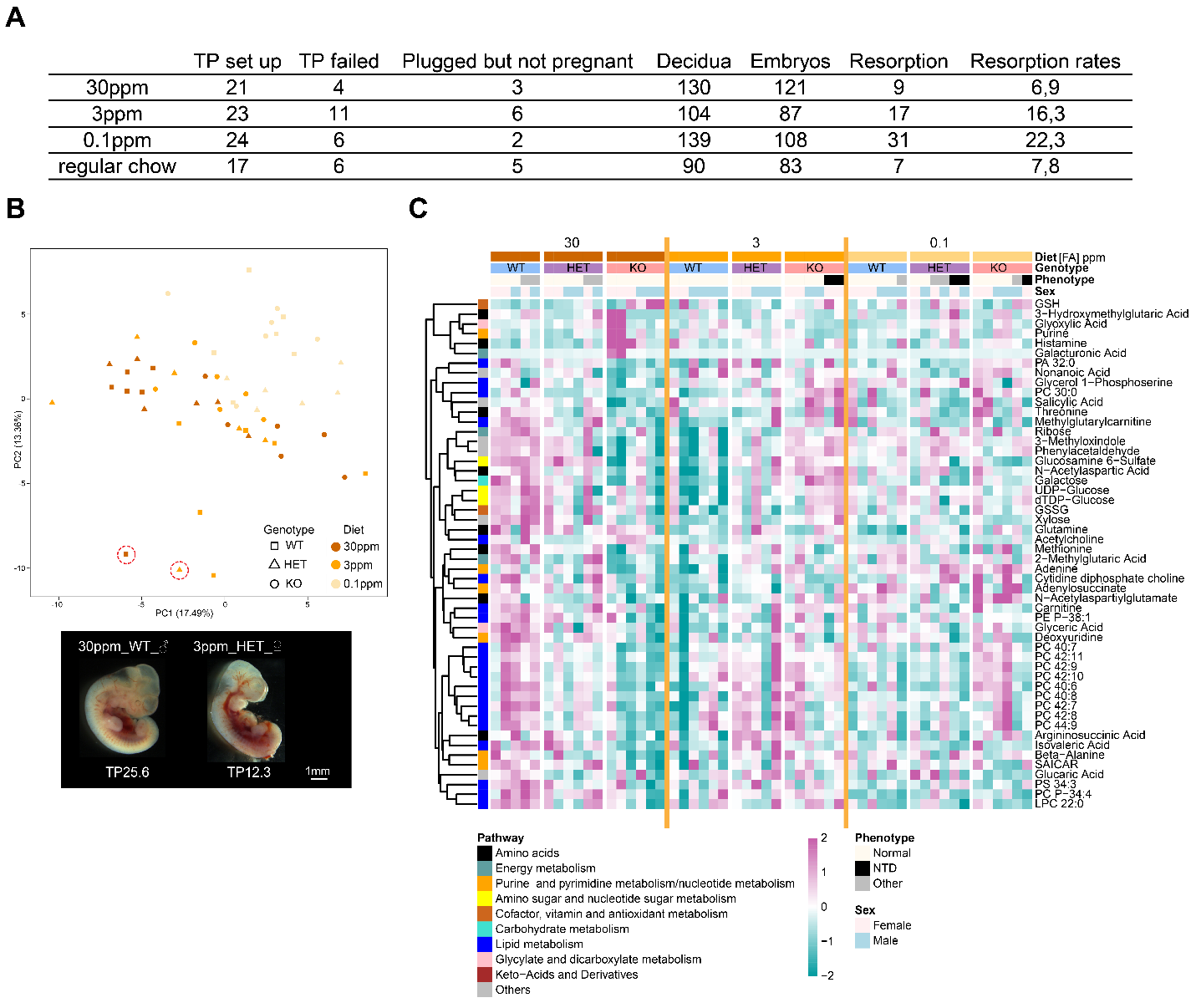

Fig. S5. Phenotypic and metabolomic changes in 129S6.Cg embryos in response to maternal dietary FA excess or depletion, interacting with the loss of *Tet1* (related to main Fig. 4).

(**A**), Summary table of timed pregnancy (TP) set up in the 129S6.Cg across all three FA-modified custom diets and regular maintenance chow. Plugged but not pregnant, indicates detection of vaginal plug, but no pregnancy at E11.5 end-point. TP failed, no successful mating based on absence of plug and pregnancy. (**B**), Principal Component Analysis of individual E11.5 whole embryo used in UPLC-MS/MS. Two outliners in red circle are excluded in downstream 2-way ANOVA interaction analysis. Embryo images of outliners are shown in the bottom panel. (**C**), Expanded heatmap of Fig. 4(F), showing individual embryo per genotype per diet group with details of phenotype and sex in the 129S6.B6-*Tet1*^tm1Koh^ strain. 2-way ANOVA interaction *P* < 0.05. n=6 per genotype per diet, except n=5 in HET 3 ppm FA and WT 30 ppm FA groups.

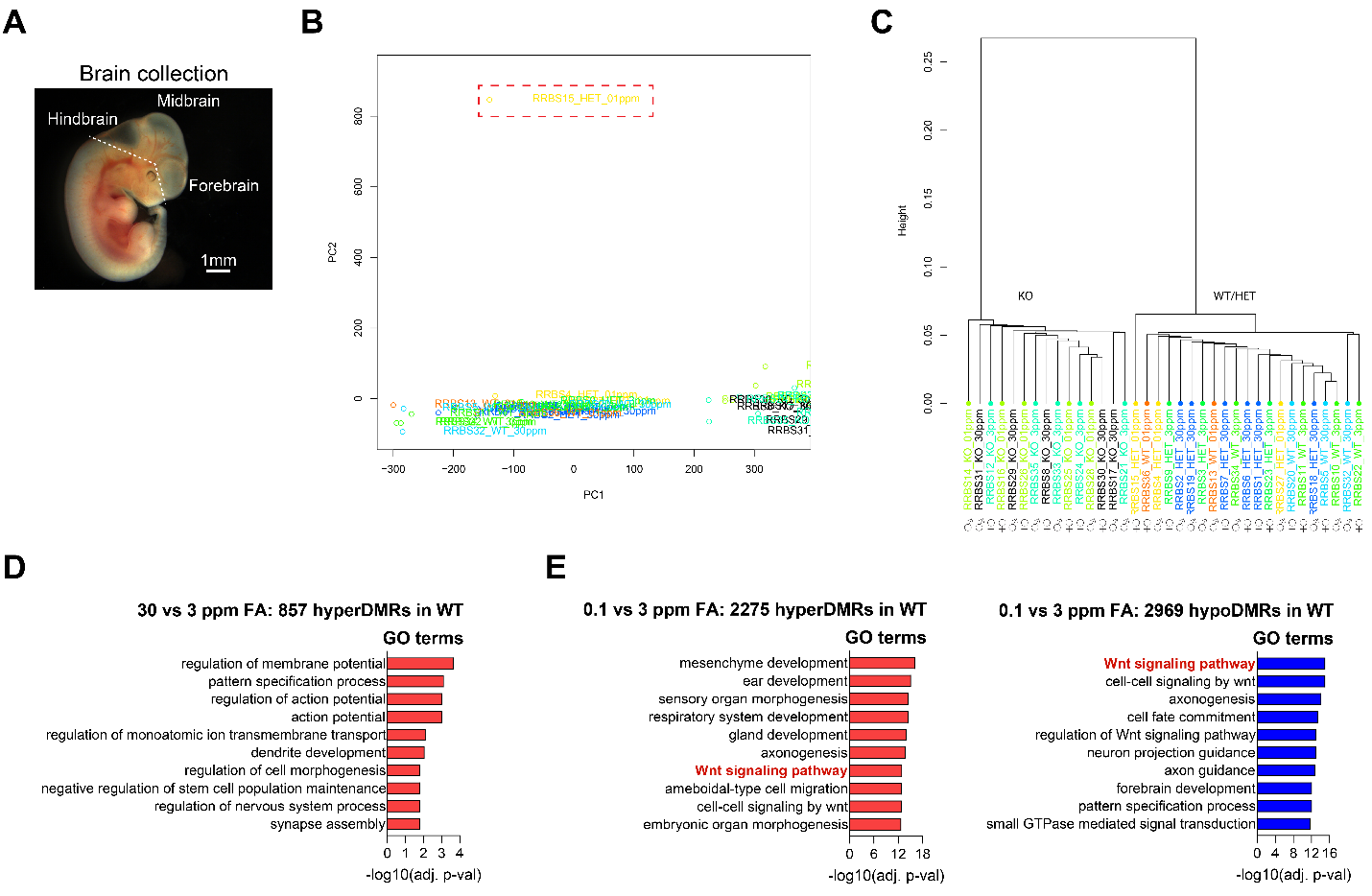

Fig. S6. RRBS analysis of 129S6.Cg E11.5 embryonic brains in response to maternal dietary FA excess or depletion, interacting with the loss of *Tet1* (related to main Fig. 5).

(**A**), Schematic diagram of E11.5 embryonic brain dissection. Plan of dissection is shown as dotted line, above which embryonic brain tissues were collected in RRBS and RNA-seq experiments. (**B**), Principal Component Analysis of individual E11.5 embryonic brains used in RRBS. One outliner was excluded in downstream analysis because of poor mapping coverage. (**C**), Hierarchical clustering plot of individual samples after exclusion of one outliner. Left, cluster of KO samples; right, cluster of WT and HET samples. n=5, KO per diet group, matched to n=3 WT and n=6 HET under 30ppm FA, n=4 WT and n=3 HET under 3ppm FA, and n=2 WT and n=3 HET under 0.1ppm FA. (**D**), GO analysis of 2275 hyperDMRs identified in WT 30 vs 3 ppm FA. (**E**), GO analysis of 2275 hyperDMRs and 2969 hypoDMRs identified in WT 0.1 vs 3 ppm FA.

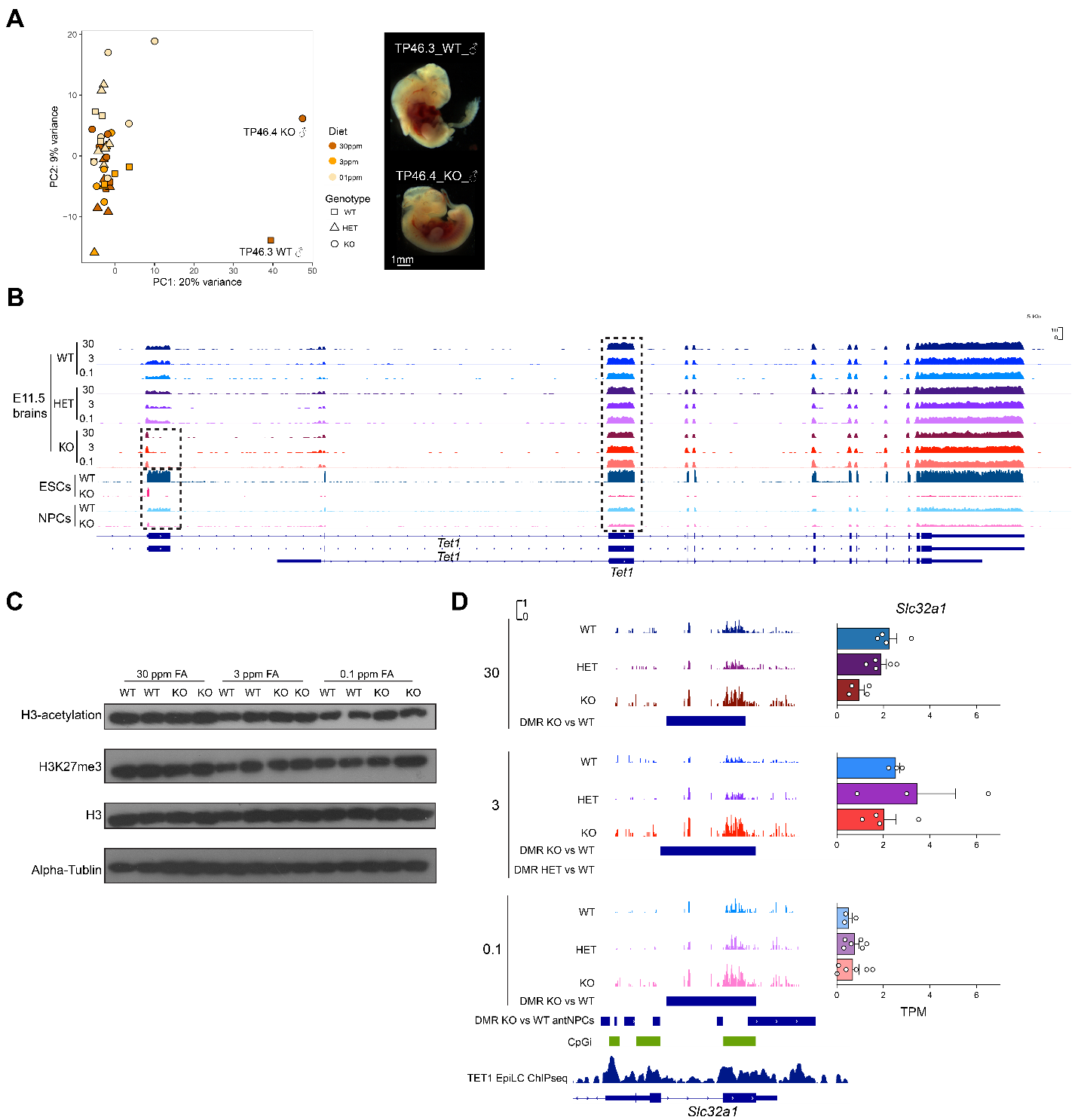

Fig. S7. RNA-seq analysis of 129S6.Cg E11.5 embryonic brains in response to maternal dietary FA excess or depletion, interacting with the loss of *Tet1* (related to main Fig. 6).

(**A**), Principal Component Analysis of individual E11.5 embryonic brain used in bulk RNA-seq. Two outliners are marked on the plot and excluded in downstream analysis. Embryo images of outliners are shown in the right panel. (**B**), Integrative Genomics Viewer (IGV) tracks of RNA-seq signals over the *Tet1* gene locus in E11.5 WT, HET, and KO embryonic brains per custom diet group, and in WT, KO ESC and differentiated NPC cell lines. Dashed box on the left denotes ablation of transcripts in the 5’ exon targeted by the knockin-knockout construct. Dashed box over a downstream exon on the right indicates expression of a short *Tet1* isoform in the E11.5 KO brains form a downstream TSS. Annotations of the embryonic full-length *Tet1* and the short (somatic) *Tet1* transcript isoforms are indicated in the bottom panel. (**C**), Western blot of pan histone 3 acetylation and H3K27me3 in KO and WT embryonic brains exposed to 30 ppm, 3 ppm and 0.1 ppm FA in the 129S6.Cg-*Tet1*^tm1Koh^ strain. ACTB, beta-actin loading control. (**D**), IGV tracks of RRBS CpG methylation levels over a DMR at *Slc32a1* in WT, HET, and KO brains per diet group (left). Locations of DMRs identified from WGBS of KO vs WT antNPCs, CpG island (CpGi) annotation and TET1 ChIP-seq signals in EpiLC, are indicated in the bottom panel. RNA-seq TPM expression of *Slc32a1* per group is shown on the right.

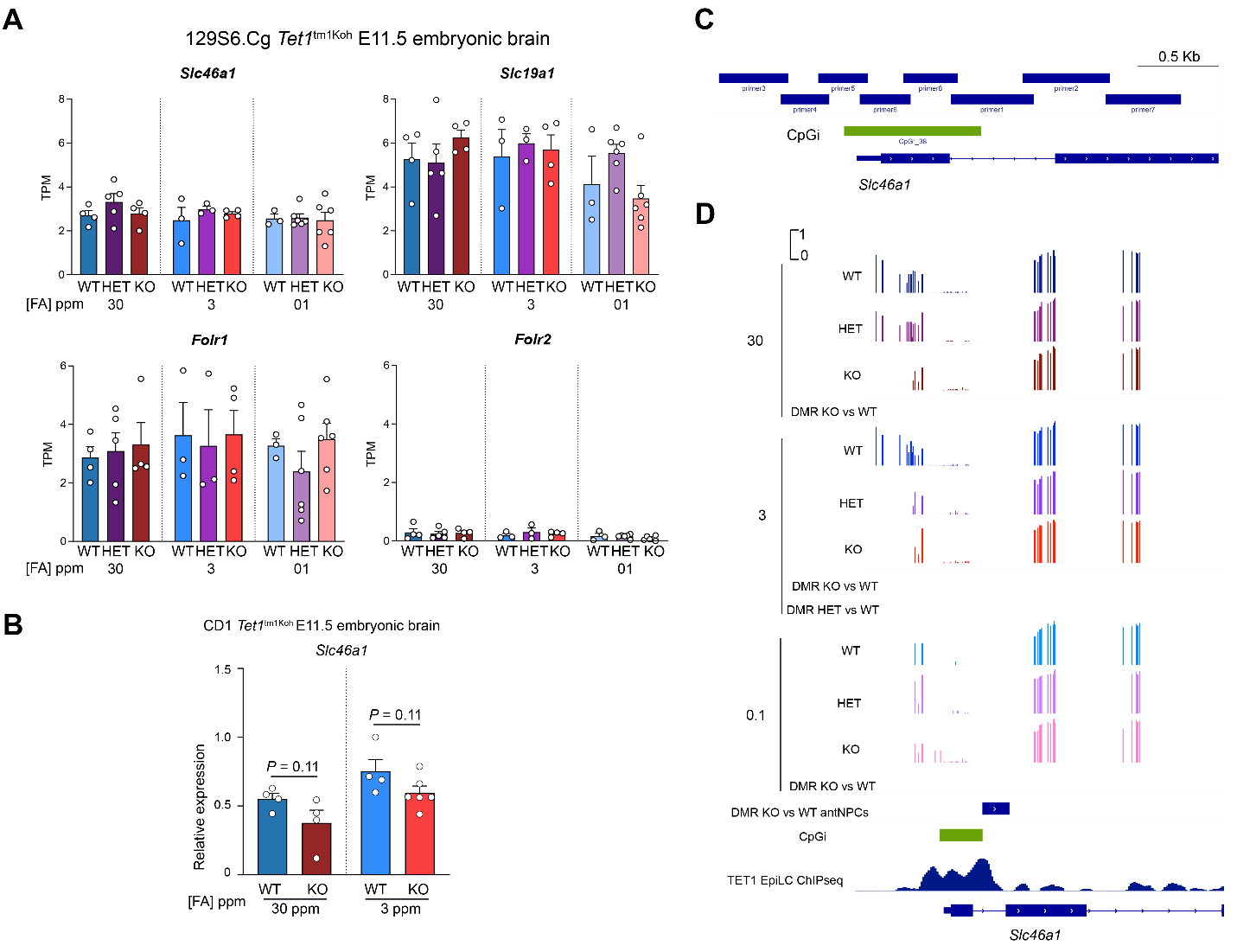

Fig. S8. Gene expression and DNA methylation analysis of folate transporters (related to main Fig. 7).

(**A**), RNA-seq gene expression of folate transporters *Slc46a1* and *Slc19a1* (top row), folate receptors *Folr1* and *Folr2* (bottom row) in 129S6.Cg E11.5 embryonic brains by *Tet1* genotype and diet group**.** (**B**), QPCR analysis of *Slc46a1* in E11.5 embryonic brains exposed to 30 ppm and 3 ppm FA in CD1-*Tet1*^tm1Koh^ mice. n = 6, KO in 3 ppm FA and n=4, all other groups. *P* values are calculated by Student’s t-test. (**C**), Overview of primer amplicons designed to cover the promoter region (~1.3 Mb) of *Slc46a1*. (**D**), IGV tracks of RRBS CpG methylation levels over the *Slc46a1* promoter region in WT, HET, and KO brains per custom diet (top), indicating a CpGi shore region with no coverage by RRBS. Locations of DMR from WGBS of KO vs WT antNPCs, CpG island (CpGi) annotation and TET1 ChIP-seq signals in EpiLC, are indicated in the bottom panel.

Table S1.

| Type | Oligo name | Sequence (5’ to 3’) |
| --- | --- | --- |
| Tet1<tm1> genotyping | Tet1tm1-Gtype-7F | TTGGCAACACCTCCAGATT |
|  | Tet1tm1-Gtype-6R | CGGATTGACCGTAATGGGATAG |
|  | Tet1tm1-Gtype-10R | GCTTTGATGTCTTCGTCTTCATC |
| Sex genotyping | SX-F | GATGATTTGAGTGGAAATGTGAGGTA |
|  | SX-R | CTTATGTTTATAGGCATGCACCATGTA |
| SNV genotyping | Cilp_SNP _Gtype_FW | GGAGAGGCTGGTGCTCACAT |
|  | Cilp_SNP _Gtype_RV | CCAGAGTTTCATGGCTGGCACA |
|  | Dapk2_SNP _Gtype_FW | CCAACATCATCACGCTGCAC |
|  | Dapk2_SNP _Gtype_Rv | CAGGGCAGAGCAAGATGAC |

Table S2.

Sample lists used for HPLC-MS/MS, RRBS, and RNA-seq on 129S6 E11.5 embryonic brains.

**HPLC-MS/MS Sample list**

| **TP** | **Group/Diet** | **Genotype** | **Phenotype** | **TS** | **Gender** | **Comments** |
| --- | --- | --- | --- | --- | --- | --- |
| TP15.6 | 30ppm+SST | WT | Indented Hindbrain | 19/20 | ♀ |  |
| TP20.8 | 30ppm+SST | WT | Normal | 19/20 | ♀ |  |
| TP29.2 | 30ppm+SST | WT | Normal | 19/20 | ♀ |  |
| TP20.6 | 30ppm+SST | WT | Hindbrain deformity | 19/20 | ♂ |  |
| TP25.6 | 30ppm+SST | WT | Normal | 19/20 | ♂ | excluded |
| TP29.5 | 30ppm+SST | WT | Normal | 19/20 | ♂ |  |
| TP20.5 | 30ppm+SST | HET | Hindbrain deformity | 19/20 | ♀ |  |
| TP11.8 | 30ppm+SST | HET | Normal | 19/20 | ♀ |  |
| TP25.3 | 30ppm+SST | HET | Normal | 19/20 | ♀ |  |
| TP26.4 | 30ppm+SST | HET | Hindbrain deformity | 19/20 | ♂ |  |
| TP11.3 | 30ppm+SST | HET | Normal | 19/20 | ♂ |  |
| TP15.7 | 30ppm+SST | HET | Normal | 19/20 | ♂ |  |
| TP11.7 | 30ppm+SST | KO | Normal | 19/20 | ♀ |  |
| TP15.4 | 30ppm+SST | KO | Normal | 19/20 | ♀ |  |
| TP20.1 | 30ppm+SST | KO | Normal | 19/20 | ♀ |  |
| TP20.2 | 30ppm+SST | KO | Normal | 19/20 | ♂ |  |
| TP26.2 | 30ppm+SST | KO | Normal | 19/20 | ♂ |  |
| TP29.1 | 30ppm+SST | KO | Normal | 19/20 | ♂ |  |
| TP18.4 | 3ppm+SST | WT | Normal | 20 | ♀ | 2♀ |
| TP24.1 | 3ppm+SST | WT | Normal | 19/20 | ♀ | 4♂ |
| TP24.2 | 3ppm+SST | WT | Normal | 19/20 | ♂ |  |
| TP27.1 | 3ppm+SST | WT | Normal | 19/20 | ♂ |  |
| TP27.2 | 3ppm+SST | WT | Normal | 19/20 | ♂ |  |
| TP33.8 | 3ppm+SST | WT | Normal | 19/20 | ♂ |  |
| TP12.3 | 3ppm+SST | HET | Delayed | 18 | ♀ | excluded |
| TP12.7 | 3ppm+SST | HET | Normal | 19/20 | ♀ |  |
| TP27.4 | 3ppm+SST | HET | Normal | 19/20 | ♀ |  |
| TP12.8 | 3ppm+SST | HET | Normal | 19/20 | ♂ |  |
| TP24.5 | 3ppm+SST | HET | Normal | 19/20 | ♂ |  |
| TP30.1 | 3ppm+SST | HET | Normal | 20 | ♂ |  |
| TP12.1 | 3ppm+SST | KO | NTD | 19/20 | ♀ | 4♀ |
| TP30.2 | 3ppm+SST | KO | NTD | 20 | ♀ | 2♂ |
| TP27.5 | 3ppm+SST | KO | Normal | 19/20 | ♀ |  |
| TP33.1 | 3ppm+SST | KO | Normal | 19/20 | ♀ |  |
| TP18.3 | 3ppm+SST | KO | Normal | 20 | ♂ |  |
| TP30.3 | 3ppm+SST | KO | Normal | 20 | ♂ |  |
| TP14.7 | 0.1ppm+SST | WT | Normal | 20 | ♀ |  |
| TP21.7 | 0.1ppm+SST | WT | Normal | 19/20 | ♀ |  |
| TP32.4 | 0.1ppm+SST | WT | Normal | 20 | ♀ |  |
| TP21.2 | 0.1ppm+SST | WT | Hindbrain deformity | 19/20 | ♂ |  |
| TP21.6 | 0.1ppm+SST | WT | Normal | 19/20 | ♂ |  |
| TP23.6 | 0.1ppm+SST | WT | Normal | 19/20 | ♂ |  |
| TP16.5 | 0.1ppm+SST | HET | Hindbrain deformity | 19 | ♀ |  |
| TP23.1 | 0.1ppm+SST | HET | NTD | 19 | ♀ |  |
| TP28.4 | 0.1ppm+SST | HET | Normal | 19 | ♀ |  |
| TP16.7 | 0.1ppm+SST | HET | NTD | 18 | ♂ |  |
| TP23.5 | 0.1ppm+SST | HET | Hindbrain deformity | 19 | ♂ |  |
| TP14.2 | 0.1ppm+SST | HET | Normal | 20 | ♂ |  |
| TP14.5 | 0.1ppm+SST | KO | Normal | 20 | ♀ |  |
| TP21.4 | 0.1ppm+SST | KO | NTD, delayed | 18/19 | ♀ |  |
| TP32.1 | 0.1ppm+SST | KO | Normal | 20 | ♀ |  |
| TP14.4 | 0.1ppm+SST | KO | Hemorrhagic | 20 | ♂ |  |
| TP16.1 | 0.1ppm+SST | KO | Normal | 19 | ♂ |  |
| TP23.3 | 0.1ppm+SST | KO | Normal | 19 | ♂ |  |

**RRBS Sample list**

| **TP** | **Group/Diet** | **Genotype** | **Phenotype** | **TS** | **Gender** | **Comments** |
| --- | --- | --- | --- | --- | --- | --- |
| 36.2 | 30ppm + SST | HET | Normal | ♀ | 19/20 |  |
| 36.3 | 30ppm + SST | WT | Normal | ♀ | 19/20 |  |
| 36.4 | 30ppm + SST | HET | Hindbrain deformity | ♀ | 19/20 |  |
| 36.7 | 30ppm + SST | KO | NTD | ♀ | 19/20 |  |
| 38.3 | 30ppm + SST | HET | Normal | ♂ | 19/20 |  |
| 38.6 | 30ppm + SST | KO | NTD | ♂ | 19/20 |  |
| 38.7 | 30ppm + SST | HET | Hindbrain deformity | ♂ | 19/20 |  |
| 41.2 | 30ppm + SST | HET | Normal | ♂ | 19/20 |  |
| 41.3 | 30ppm + SST | WT | Normal | ♀ | 19/20 |  |
| 41.4 | 30ppm + SST | KO | Hindbrain deformity | ♂ | 19/20 |  |
| 41.8 | 30ppm + SST | KO | Normal | ♀ | 19 |  |
| 43.4 | 30ppm + SST | KO | Normal | ♂ | 19/20 |  |
| 43.7 | 30ppm + SST | WT | Normal | ♂ | 19/20 |  |
| 40.1 | 3ppm + SST | HET | Normal | ♀ | 19 |  |
| 40.2 | 3ppm + SST | WT | Normal | ♂ | 19/20 |  |
| 40.3 | 3ppm + SST | WT | Normal | ♀ | 19/20 |  |
| 40.5 | 3ppm + SST | KO | Normal | ♀ | 19 |  |
| 40.10 | 3ppm + SST | KO | Normal | ♂ | 19 |  |
| 42.1 | 3ppm + SST | HET | Normal | ♂ | 19/20 |  |
| 45.3 | 3ppm + SST | WT | Normal | ♀ | 19/20 |  |
| 45.6 | 3ppm + SST | HET | Normal | ♀ | 19/20 |  |
| 45.8 | 3ppm + SST | KO | Normal | ♀ | 19/20 |  |
| 42.9 | 3ppm + SST | KO | Hindbrain deformity | ♂ | 19/20 |  |
| 57.3 | 3ppm + SST | WT | Normal | ♂ | 19/20 |  |
| 57.5 | 3ppm + SST | KO | NTD | ♂ | 19/20 |  |
| 34.1 | 0.1ppm + SST | HET | Normal | ♂ | 19/20 |  |
| 34.3 | 0.1ppm + SST | WT | Normal | ♂ | 19/20 |  |
| 34.5 | 0.1ppm + SST | KO | Normal | ♂ | 19/20 |  |
| 35.2 | 0.1ppm + SST | HET | Normal | ♀ | 19/20 | excluded |
| 35.4 | 0.1ppm + SST | KO | Hindbrain deformity | ♀ | 19/20 |  |
| 50.1 | 0.1ppm + SST | WT | Hindbrain deformity | ♀ | 19/20 |  |
| 50.3 | 0.1ppm + SST | KO | NTD | ♀ | 19/20 |  |
| 50.5 | 0.1ppm + SST | KO | NTD | ♀ | 19/20 |  |
| 54.7 | 0.1ppm + SST | HET | Normal | ♂ | 19 |  |
| 54.8 | 0.1ppm + SST | KO | Normal | ♂ | 19 |  |

**RNA-seq Sample list**

| **TP** | **Group/Diet** | **Genotype** | **Phenotype** | **TS** | **Gender** | **Comments** | **RIN** |
| --- | --- | --- | --- | --- | --- | --- | --- |
| 38.1 | 30ppm + SST | WT | Normal | ♀ | 19/20 |  | 10 |
| 38.4 | 30ppm + SST | HET | Hindbrain deformity | ♀ | 19/20 |  | 10 |
| 43.2 | 30ppm + SST | WT | Normal | ♀ | 19/20 |  | 10 |
| 43.3 | 30ppm + SST | HET | Normal | ♀ | 19/20 |  | 10 |
| 43.6 | 30ppm + SST | KO | Normal | ♂ | 19/20 |  | 10 |
| 46.3 | 30ppm + SST | WT | Grossly malformed | ♂ | 19/20 | excluded | 9,2 |
| 46.4 | 30ppm + SST | KO | Grossly malformed | ♂ | 19/20 | excluded | 9,1 |
| 46.7 | 30ppm + SST | HET | Normal | ♂ | 19/20 |  | 10 |
| 46.8 | 30ppm + SST | HET | Hindbrain deformity | ♂ | 19/20 |  | 10 |
| 47.1 | 30ppm + SST | WT | Normal | ♂ | 19/20 |  | 10 |
| 47.3 | 30ppm + SST | KO | NTD | ♀ | 19 |  | 10 |
| 47.5 | 30ppm + SST | HET | Normal | ♀ | 19/20 |  | 10 |
| 47.6 | 30ppm + SST | KO | Normal | ♀ | 19/20 |  | 10 |
| 49.1 | 30ppm + SST | WT | Normal | ♀ | 19/20 |  | 10 |
| 49.2 | 30ppm + SST | KO | Normal | ♀ | 19/20 |  | 10 |
| 45.1 | 3ppm + SST | KO | Normal | ♂ | 19/20 |  | 10 |
| 45.4 | 3ppm + SST | HET | Normal | ♂ | 19/20 |  | 10 |
| 51.2 | 3ppm + SST | HET | Normal | ♀ | 20 |  | 10 |
| 51.6 | 3ppm + SST | KO | Normal | ♀ | 20 |  | 10 |
| 51.9 | 3ppm + SST | WT | Normal | ♀ | 20 |  | 10 |
| 53.1 | 3ppm + SST | KO | Normal | ♂ | 19/20 |  | 10 |
| 53.2 | 3ppm + SST | WT | Normal | ♂ | 19/20 |  | 10 |
| 53.4 | 3ppm + SST | HET | Normal | ♂ | 19/20 |  | 10 |
| 57.6 | 3ppm + SST | KO | NTD | ♂ | 19/20 |  | 10 |
| 57.8 | 3ppm + SST | WT | Normal | ♂ | 19/20 |  | 10 |
| 34.2 | 0.1ppm + SST | HET | Normal | ♀ | 19/20 |  | 10 |
| 34.4 | 0.1ppm + SST | HET | Hindbrain deformity | ♀ | 19/20 |  | 9,9 |
| 34.7 | 0.1ppm + SST | KO | Normal | ♀ | 19 |  | 10 |
| 35.1 | 0.1ppm + SST | HET | Normal | ♂ | 19/20 |  | 10 |
| 35.5 | 0.1ppm + SST | KO | Normal | ♂ | 19/20 |  | 10 |
| 48.1 | 0.1ppm + SST | KO | Normal | ♂ | 19/20 |  | 10 |
| 48.5 | 0.1ppm + SST | HET | Normal | ♂ | 19/20 |  | 10 |
| 50.6 | 0.1ppm + SST | KO | NTD | ♀ | 19/20 |  | 9,6 |
| 52.1 | 0.1ppm + SST | WT | Normal | ♀ | 19 |  | 9,9 |
| 52.2 | 0.1ppm + SST | WT | Normal | ♂ | 19 |  | 10 |
| 52.3 | 0.1ppm + SST | WT | Hindbrain deformity | ♀ | 19 |  | 9,9 |
| 52.5 | 0.1ppm + SST | KO | Delay | ♂ | 18 |  | 10 |
| 54.1 | 0.1ppm + SST | KO | Normal | ♂ | 19/20 |  | 9,7 |
| 54.2 | 0.1ppm + SST | HET | Normal | ♂ | 19/20 |  | 10 |
| 54.3 | 0.1ppm + SST | HET | Hindbrain deformity | ♂ | 19/20 |  | 9,8 |
